# Ancient and ongoing hybridization in the *Oreochromis* cichlid fishes

**DOI:** 10.1101/2023.05.19.541459

**Authors:** Adam G Ciezarek, Tarang K Mehta, Angela Man, Antonia GP Ford, Geraldine Dorcas Kavembe, Nasser Kasozi, Benjamin P Ngatunga, Asilatu H Shechonge, Rashid Tamatamah, Dorothy Wanja Nyingi, Avner Cnaani, Federica Di Palma, George F Turner, Martin J Genner, Wilfried Haerty

## Abstract

Hybridization may enable adaptive diversification by generating unique genetic combinations when hybrid lineages are faced with ecological opportunity. Conversely, hybridization with exotic species may be detrimental to native biodiversity, by leading to homogenisation and the loss of important genetic material associated with local adaptation. Here we studied both ancient and contemporary hybridization in cichlid fishes of the genus *Oreochromis* (tilapia), which are among the most important fish for global aquaculture. We use whole genome resequencing of 575 individuals from 23 species, focussing on Tanzania, a natural hotspot of tilapia diversity, and a country where hybridization between exotic and native species in the natural environment has been previously reported. We reconstruct the first genome-scale phylogeny of the genus and reveal prevalent ancient gene flow across the *Oreochromis* phylogeny. This introgression has not led to large-scale adaptive radiation as seen in other cichlid lineages. We identify multiple cases of contemporary hybridization between native and introduced species in the wild, linked to the use of non-native species in aquaculture improvement and stocking for capture fisheries. Our study shows how ancient hybridization contributed to modern tilapia diversity, but is now a threat to both the genetic integrity of wild populations and the long-term prospects of the global tilapia aquaculture industry.

## Introduction

Global aquaculture production has increased dramatically in recent decades, and is projected to take a key role in meeting the food demands of a growing human population, whilst alleviating nutritional poverty (Ahmed et al., 2019; Hall et al., 2017; Naylor et al., 2021). Tilapia of the genus *Oreochromis* (Cichlidae: Oreochromini) are a group of cichlid fish native to Africa and the Middle East, and are now the second largest group (by tonnage produced) of any aquaculture fish globally, following carp (Cyprinidae). *Oreochromis* production is dominated by a single species, the Nile tilapia, *Oreochromis niloticus*, which accounted for 4.5 out of the total 5.5 million tonnes of tilapia produced in 2020, the third most of any fish species globally. The remaining ∼1 million tonnes are only accurately recorded to the genus level (FAO, 2022). The success of Nile tilapia has been attributed to high growth rates as well as tolerance of a wide range of environmental conditions, such as temperature, salinity, pH and low dissolved oxygen enable (El-Sayed & Fitzsimmons, 2023). In principle, the use of wild relatives of the Nile tilapia in breeding programmes could be used to further enhance production (Lind et al., 2012). However, despite this importance for food security, many of the 37 described *Oreochromis* species (Ford et al., 2019) are poorly known.

Although *Oreochromis* tilapia are widespread across Africa, subject to major capture fisheries, and key aquaculture species, considerably more attention has been given to the evolution of haplochromine cichlid systems (Joyce et al., 2011; Salzburger, 2018; Seehausen, 2006; Sturmbauer, 1998). This focus has been due to the propensity of haplochromines to form expansive lacustrine adaptive radiations, characterised by rapid speciation and extensive ecomorphological diversification (Malinsky et al., 2018; McGee et al., 2020). This occurred despite low levels of genetic diversity, a high proportion of which is shared between species due to incomplete lineage sorting and widespread interspecific introgression (Svardal et al., 2021). It has been proposed that extensive lacustrine radiations of haplochromine cichlids have been facilitated by the introgression arising from hybridization of ancestral lineages (Meier et al., 2017; Svardal et al., 2020). This may fuel differentiation and diversification by supplying novel combinations of genetic variants (Singhal et al., 2021), and sharing of ancient alleles that have already been filtered by selection (Marques et al., 2019). By contrast, ongoing or recurring hybridization may decrease differentiation and homogenise gene pools, therefore reducing the propensity of a lineage to radiate.

In contrast to the haplochromines, extensive radiation has not taken place in the *Oreochromis* (Seehausen, 2006). Most *Oreochromis* species occupy rivers and are allopatric to apparent sister taxa (Ford et al., 2019; Trewavas, 1983); the only known *Oreochromis* lacustrine radiations are small and limited to Lake Malawi and Lake Natron (3 species each). To date, comprehensive phylogenomic examinations of ancestral introgression occurring during the evolution of *Oreochromis* are lacking, and therefore it is currently not clear what role ancestral introgression has played in *Oreochromis* diversification.

Given the high productivity of some *Oreochromis* species in aquaculture and capture fisheries, they have been widely introduced across tropical and subtropical freshwaters globally (Canonico et al., 2005; Lévêque, 1995). In Africa, introductions to the natural environment have resulted in widely documented contemporary hybridization (Blackwell et al., 2020; Bradbeer et al., 2019; Champneys et al., 2021; Ciezarek et al., 2022; Shechonge et al., 2018). The risk of hybridisation between native and exotic *Oreochromis* species is likely to increase with the projected growth of aquaculture across Africa (Prabu et al., 2019). Detailed analysis of the extent of hybridization between co-occurring tilapia species will therefore be valuable for identifying species and populations of conservation concern.

This study focuses on the genomic implications on ancestral and contemporary introgression in *Oreochromis*, focusing on wild populations in Tanzania. The country is a hotspot for diversity of the genus with at least 21 species present, including a minimum of three species that have been widely introduced outside their native range, namely *O. niloticus*, *O. leucostictus* and *O. esculentus* (Shechonge et al., 2019). To reconstruct the phylogenetic relationships between species, and to quantify ancestral and modern hybridization, we present a genome-wide resequencing dataset of the genus *Oreochromis*. We provide a first phylogenomic analysis of the group and show that *Oreochromis* is typified by a high degree of ancestral introgression. We also identify a species of hybrid origin, the Lake Kiungululu tilapia, *O. chungruruensis*. We further find that anthropogenic activities have resulted in widespread recent hybridization, highlighting a potential risk to the conservation of species and their genetic diversity.

## Materials and methods

### Sampling and sequencing

We sampled and generated sequencing data for 433 individuals, and combined this with previously published data from 142 individuals (Table S1) from across southern and eastern Africa (Figure 1). Samples were from 65 locations, and 21 drainage basins across Tanzania, as well as South Africa, Kenya, Angola, Malawi, Mozambique and Uganda. Individuals were morphologically identified to species using information outlined in (Genner et al., 2018). Individuals were classified as hybrids if they had characteristics intermediate between different species. Our final dataset comprised of 25 species, including the outgroups *Maylandia zebra* and *Sarotherodon galilaeus* and 23 out of the 37 species of *Oreochromis* currently recognised, including *Oreochromis (Alcolapia) grahami* (Fricke et al., 2023). Fish specimens were mostly obtained from local fishers or through experimental fishing surveys. Live fish were euthanised with an overdose of anaesthetic (MS-222). Samples for subsequent DNA extraction were collected by clipping off the right pectoral fin, which was placed in a labelled vial of ethanol. Whole voucher specimens were retained, with each individual pinned on a styrofoam board, photographed and preserved in formalin or ethanol.

**Figure 1.**
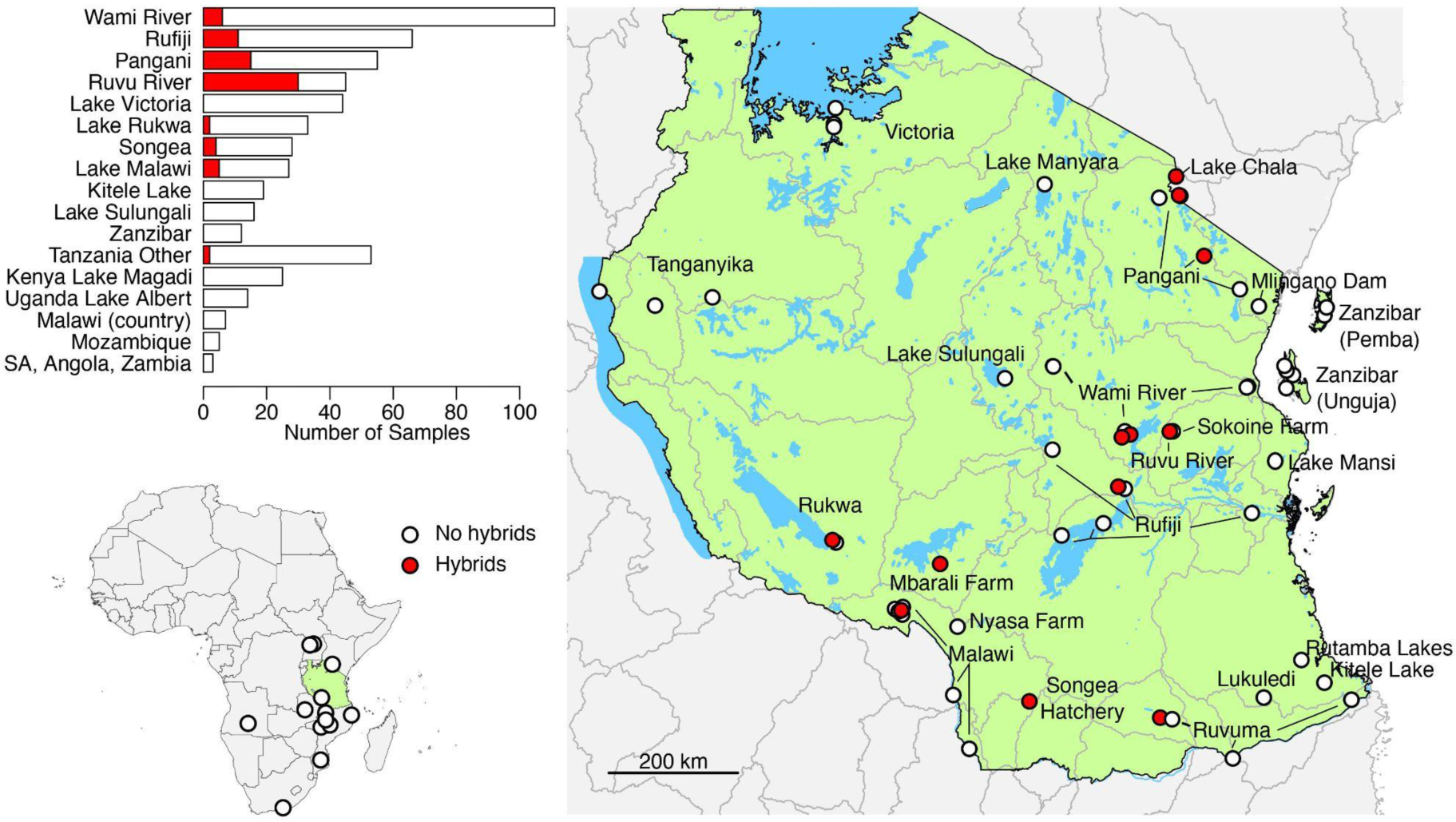
Locations of the *Oreochromis* sampled across Africa (bottom left) and, within Tanzania (right). Sample sites where contemporary hybrids were identified from genomic data are highlighted in red. Key drainage basins are labelled. The bar chart (top left) shows the total number of specimens (contemporary hybrids in red, non-hybrids in white) located per drainage basin within Tanzania, for all basins with more than ten samples, or country outside of Tanzania. All basins within Tanzania with fewer than ten specimens sampled are grouped into “Tanzania Other”. Abbreviation: SA - South Africa. Geographic coordinates of sampling locations are shown in Table S1.

DNA was extracted from fin clips using a PureLink Genomic DNA extraction kit (Life Technologies). Sequencing was conducted using either PCR-free library preparation with Illumina HiSeq 2500 (250bp paired-end), automated KAPA DNA library preparation with Illumina HiSeq 2500 (125bp paired-end), LITE library preparation with Illumina HiSeq 4000 (150bp paired-end) or LITE library preparation with NovaSeq S4 (150bp paired-end).

### SNP calling

SNPs were called against previously published near chromosome-level assemblies of *M. zebra* (GCF_000238955.4) and *O. niloticus* (GCF_001858045.2) (Conte et al., 2019). Raw reads were checked for adapter sequence contamination using BBMerge, within BBMap (v38.06) (Bushnell, 2014). Where necessary, reads were trimmed for adapters using fastp (v0.20.0) (Chen et al., 2018), with quality-trimming disabled. Reads were then mapped separately against both assemblies using the mem function in bwa (v0.30.0) (Li & Durbin, 2009), with raw mappings sorted by coordinate and having mate coordinates and insert size fields added using samtools (v1.9) (Li et al., 2009). Joint genotyping was then carried out on all 575 samples using bcftools (v1.10.2) (Li et al., 2009). First, bcftools mpileup was used with a minimum base and mapping qualities of 30. Multi-allelic variant calling was then carried out using bcftools call. Variants were filtered if they were within three base pairs of any other variant, if variant quality score was less than 30, if depth at a given site (across all samples) was less than 500 or greater than 9000, or if minor allele count was less than three.

### Backbone phylogeny reconstruction

To construct a species-level phylogeny, we identified a panel of 91 reference individuals that were unlikely to be recent hybrids (Table S1). These individuals were identified using morphology, evidence of monophyly in a full genome neighbour-joining tree of individuals not classified morphologically as hybrid, and evidence of monophyly in a mitochondrial genome neighbour-joining tree of those individuals (Supplementary Data 1). Using this reference panel, separate phylogenetic trees were inferred for each of the *M. zebra* and *O. niloticus* mapped datasets. Trees were constructed using both a maximum-likelihood SNP-based concatenation tree, inferred using IQ-TREE (v1.6.12) (Nguyen et al., 2015), as well as a multi-species coalescent summary-based tree, inferred using ASTRAL (v5.4.12) (Rabiee et al., 2019; Zhang et al., 2018), based on trees inferred from recombination-free 10kb windows across the genome with IQ-TREE (Supplementary Methods). Robinson-Foulds (RF) distances (Robinson & Foulds, 1981) were calculated between each 10kb-window tree and the inferred ASTRAL species tree using the ETE 3 python package (Huerta-Cepas et al., 2016), in order to further quantify how widespread phylogenetic discordance was. For species where the inferred phylogenetic trees raised taxonomic questions, such as a lack of monophyly of all individuals assigned to a species, exploratory Principal Component Analysis (PCA) was carried out on all individuals of the relevant species, but excluding those inferred to be a contemporary hybrid (see below), using plink (v2.0.0) (Purcell et al., 2007). The PCAs on each subset of individuals used all biallelic SNPs found with a minor allele count of at least 3 in this subset of individuals, and SNPs were pruned for linkage disequilibrium (r^2^>0.6) over 20kb windows using bcftools.

### Assessment of ancestral introgression

Ancestral introgression between species was assessed using the panel of 91 reference individuals, and both the *M. zebra* and *O. niloticus* mapping datasets. We used both SNP-based D statistics (Green et al., 2010), as well as *f*-branch statistics (Malinsky et al., 2018), calculated using Dsuite (v0.4) (Malinsky et al., 2021), and genome-wide 10kb window phylogenetic-tree-based analyses. These tree-based analyses included the branch-length tests (BLT) and discordant count tests (DCT), using the scripts provided by Suvorov et al., (2022), and chi-squared tests of discordant topology frequencies inferred using IQ-TREE (Minh et al., 2020; Suvorov et al., 2022) (Supplementary Methods). To test whether introgression was more likely between species sharing a drainage basin or between closely or more distantly related species, for each species we identified the species counterpart with the highest *f*-branch (i.e. the highest degree of gene flow). We then inferred whether these species pairs were more likely to have shared modern drainage basins than expected by chance using a randomised permutation test. Similarly, we identified for each species whether their high *f-*branch was more or less phylogenetically related than the average of all the species compared (z <= -2 for less phylogenetically distant, z >= 2 for more phylogenetically distant). We then tested whether we observed more species with significantly phylogenetically similar or dissimilar counterparts than expected by chance using a permutation test.

### Analysis of putative introgression events

Putative introgression events, identified by the *f-*branch analysis, were further investigated to identify introgressed genomic regions. This was carried out using a novel statistic (Dwt; Dweighted_topo), derived from topology weightings calculated from ‘topology weighting by iterative sampling of subtrees’ (Twisst) (Martin & Van Belleghem, 2017), computed along the 22 linkage groups of the *M. zebra* reference assembly (Supplementary Methods). Dwt is designed to highlight putatively introgressed regions. Dwt = 0 indicates regions where the species tree (sptree) has the highest weighting. Dwt > 0 indicates where disc1 (the putative introgression topology) is most heavily weighted, with 1 indicating it is fully supported (no weighting for either of the other topology). Similarly, Dwt < 0 indicates where disc2 (the other alternate topology) is most heavily weighted. 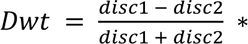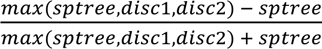.

Putative introgression events where the Twisst analysis showed no clear excess of either the species-tree topology or hybridization topology were further investigated to identify signals of hybrid speciation. Following Olave et al. (2022), we used ADMIXTURE analyses of the relevant species at both K=2 and K=3, as well as assessment of species-diagnostic SNPs. Putative introgression dates were also estimated based on estimates of Dxy and π in phylogenetically identified introgressed regions of the genome, calculated using genomics_general (https://github.com/simonhmartin/genomics_general) (see Supplementary Methods).

### Contemporary hybridization

The 486 individuals that were not among the reference panel were tested for evidence of recent hybridization. We carried out separate analyses in each sampled water drainage basin within Tanzania. For each drainage basin, biallelic SNPs were extracted for each individual sampled, alongside reference individuals for all the taxa recorded in the water body by Shechonge et al. (2019). SNPs with a minor allele count of at least 3 were retained, and SNPs were pruned for linkage disequilibrium (r^2^>0.6) over 20kb windows using bcftools. For each drainage basin, a supervised ADMIXTURE (Alexander et al., 2009) analysis was carried out only involving the relevant species or populations, with a K value equal to the number of locally recorded taxa for the ADMIXTURE analysis, and each reference species assigned to their known group. Test individuals were assigned a hybrid status if they had multiple ancestry components where the lower end of the standard error was at least 0.1. This cut-off was used to identify up to second generation backcrosses. We further confirmed that all of the reference individuals of the same species were tightly clustered using PCAs performed in plink, and comprised a monophyletic group with no long branch lengths, in phylogenetic trees inferred with IQ-TREE (v2.0) with automated model detection, 5 independent runs and 1,000 rapid bootstraps. These analyses were carried out with the same SNP set as the ADMIXTURE runs, except it was further filtered for sites where at least one individual had each of the homozygous reference and alternate alleles for the IQ-TREE analysis. We further characterised contemporary hybridization using panels of species-diagnostic SNPs (Supplementary Methods).

## Results

Mapping, phylogenomic and ancestral introgression analyses were carried out against both the *O. niloticus* and *M. zebra* reference assemblies separately. Results were similar and conclusions identical (Supplementary Materials), therefore only the *M. zebra* reference analyses are reported below.

### Sampling, sequencing, read mapping and SNP calling of 23 species

An average of 41.3 million paired-end reads were sequenced for the 433 individuals (range 10.4 - 106 million) (Figure 1). Mapped reads from all 575 individuals (including the previously published data for 142 individuals) had average depth of 6.3 (range 1.7-18.6), and average properly paired mapping of 75% (range 45-89%). In total 55,949,298 filtered SNPs were called. Full mitochondrial genomes of at least 10,000bp were successfully assembled for 437 out of the 450 individuals not morphologically identified as hybrid, with an average length of 16,541bp (range 11,048-17,126bp).

### Phylogenetic inference

A total of 1,692,136 non-coding SNPs were used for the maximum-likelihood phylogenetic inference with 15,310 recombination-free 10kb windows used for ASTRAL. Most of the taxonomic species produced monophyletic groups, and therefore collapsed to a single node. Those that were not monophyletic were instead collapsed to the population level (Figure 2a). There were some inconsistencies between the ASTRAL and maximum-likelihood trees, particularly with regard to the phylogenetic placement of *O. tanganicae, O. placidus placidus* and *O. korogwe* (Figure S2-5). The ASTRAL results also suggest *O. hunteri* to be a monophyletic species, whereas the ML trees do not (Figure S2-5). All following analyses are reported using the ASTRAL species tree, given that unlike the maximum-likelihood tree it accounts for the likely widespread incomplete lineage sorting in the dataset. This is indicated by the high levels of phylogenetic discordance shown by high Robinson-Foulds (RF) distances between each 10kb window tree and the species tree. No 10kb window tree had a RF distance from the species tree of less than 0.08 (Figure S1).

**Figure 2.**
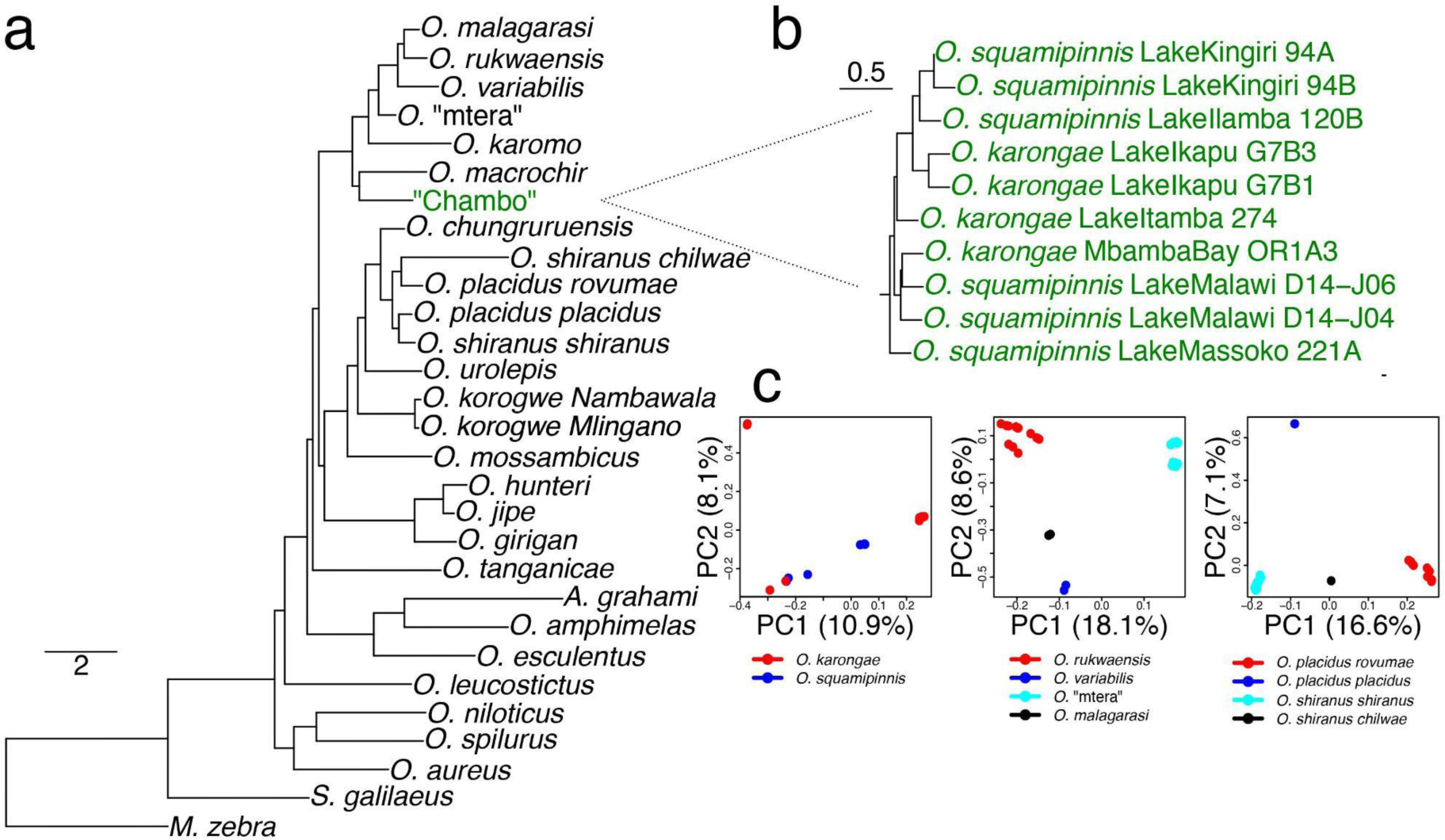
a) Phylogenetic tree inferred using ASTRAL from the *M. zebra* mapping dataset, with conspecific individuals collapsed. Branch lengths are in coalescent units. All nodes had posterior probability of 1.0. b) Phylogenetic tree inferred using ASTRAL from the *M. zebra* mapping dataset, without species collapsed, highlighting only the individuals from the Lake Malawi “Chambo” assemblage of *O. squamipinnis* and *O. karongae*. c) PCA analyses of the major taxonomic questions raised by this phylogenetic tree, showing the “Chambo” group (left), *O. rukwaensis* being distinct from *O.* “mtera” (middle), and subspecies of *O. placidus* and *O. shiranus* not supporting current classification (right).

There were instances of taxonomic species where individuals from different populations did not form monophyletic groups (Figure 2a, Figure S2), and PCA analyses further confirmed a lack of clustering of taxonomically identified species (Figure 2c). *O. rukwaensis* individuals from the Rukwa drainage basin were resolved as a clade but were phylogenetically distinct from individuals collected from the Mtera Reservoir, which had provisionally been identified as *O. rukwaensis* based on morphology (Shechonge et al., 2019). Here, we refer to this population as *O. “*mtera*”*. We found no evidence of a taxonomic split between *O. shiranus* and *O. placidus*, and instead *O. shiranus shiranus* was resolved as sister to *O. placidus placidus*, whereas *O. shiranus chilwae* is sister to *O. placidus rovumae*. Similarly, the analysed “Chambo” species, *O. squamipinnis* and *O. karongae*, are not reciprocally monophyletic in our anlayses (Figure 2b,c).

### Widespread ancestral introgression

There was consistently a high degree of ancestral introgression across the *Oreochromis* phylogeny evident in our analyses. Using the genome-wide phylogenetic trees constructed from 10kb windows, 44,030 out of the 113,564 trios tested with the blt and dct tests showed significant deviations from expected values under incomplete lineage sorting alone, consistent with introgression. Similarly, the frequencies of 10kb window-tree discordant topologies indicated introgression at 42/89 nodes in the ASTRAL tree. For the SNP-based D statistics, 2,658 out of the 2,925 tested trios showed significantly higher levels of allele sharing between non-sister taxa than expected, with Dp values indicating a range of introgression proportions ranging from 0-0.45 (Table S2; for this analysis, individuals were pooled into populations or species, hence the lower number of tested trios than the blt and dct analyses).

F-branch statistics were calculated between each branch in the tree and each tip, revealing a few branches which appeared to exhibit relatively large amounts of gene flow (Figure 3a). Notably high *f*-branch values were recorded associated with two of the taxonomic uncertainties highlighted above; namely the *O. shiranus-O. placidus* group (especially involving *O. shiranus chilwae*) and between *O. rukwaensis* and *O. “mtera”* (Figure 3a).

**Figure 3.**
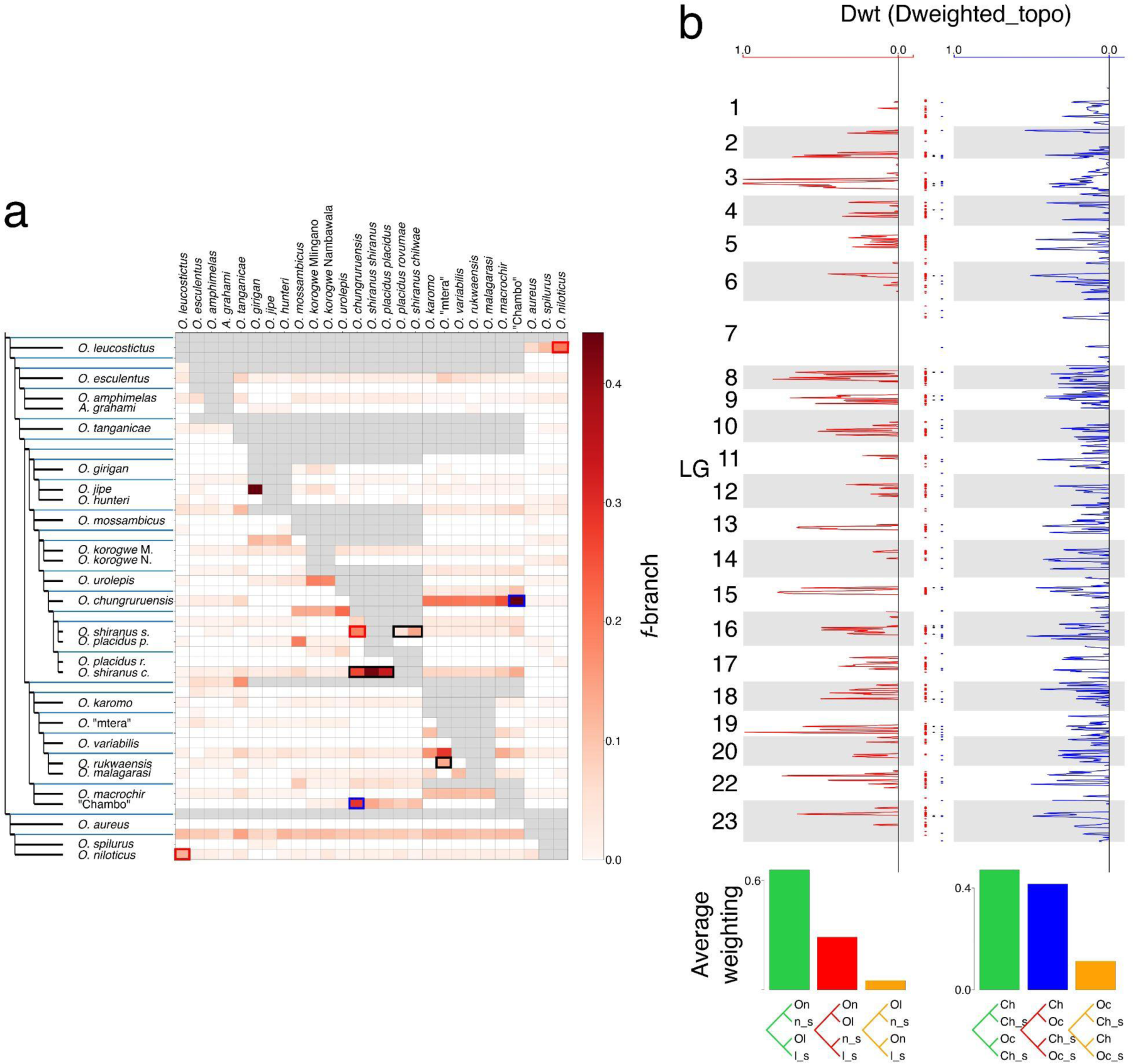
a) Heatmap of the *f*-branch values calculated between each node (y axis) and tip (x axis) of the *M. zebra* mapping ASTRAL tree. Boxes indicating the introgression from “Chambo” into *O. chungruruensis* are highlighted in blue, and between *O. leucostictus* and *O. niloticus* are highlighted in red. Boxes indicating introgression amongst the clades associated with taxonomic uncertainties are highlighted in black. b) Normalised Dwt statistics across the linkage groups showing regions with a stronger weighting for the introgression tree (left of the solid line), where *O. leucostictus* and *O. niloticus* are sister (left; red), and where “Chambo” and *O. chungruruensis* are sister (right; blue). Dots represent stretches of the genome >=50kb where the four Twisst analyses consistently have Dwt = 1.0 showing introgression between *O. leucostictus* and *O. niloticus* (red), *O. squamipinnis* and *O. chungruruensis* (blue), and overlap between both comparisons (black). Abbreviations: On - *O. niloticus*, Ol - *O. leucostictus*, On_s - *O. spilurus* + *O. aureus*; Ol_s - *O. variabilis* + *O. esculentus* + *A. grahami,* Ch - “Chambo”, Oc - *O. chungrurensis*, Ch_s - *O. rukwaensis* + *O. variabilis* + *O. macrochir*, Oc_s - *O. placidus rovumae* + *O. korogwe* Mlingano + *O. mossambicus*).

We identified the counterpart for each species which had the highest *f*-branch value. These species pairs occupied the same drainage basin more often than expected by chance (*p*=0.0002). This was the case for seven species (compared to simulated range of 0-8 species): *O. leucostictus* / *O. niloticus*, *O. girigan* / *O. korogwe* Mlingano, *O. karomo* / *O. tanganicae*, “Chambo” / *O. chungrurensis*, *O. jipe* / *O. girigan*, *O. chungruruensis* / “Chambo”, and *O. shiranus shiranus* / *O. chungruruensis*. Three out of these seven species pairs were related to a single putative hybrid speciation event of *O. chungruruensis* (see results below). After removing these species from the analysis, shared drainage basin less strongly predicted the *f*-branch values (*p*=0.03; 4 species, simulated range 0-7). Similarly, the species pairs with evidence of introgression were more likely to be more closely related than the mean phylogenetic distance between each species (Z-score <= -2) than expected by chance (*p*=0.0004). This was the case for four species pairs (compared to simulated range 0-4 species): *O. jipe* with *O. girigan*, *O. amphimelas* with *O. esculentus*, *O. rukwaensis* with *O.* “mtera” and *O. malagarasi* with *O. variabilis*.

Based on the results of *f-*branch analyses and Dp values, we focused on ancestral introgression between *O. niloticus* and *O. leucostictus* and between *O. chungruruensis* and the “Chambo” group. These were the species pairs indicating the highest proportions of introgression according to Dp (Table S2), as well as highest *f-*branch values between non-sister or closely related branches (Figure S6). Four analyses were carried out to assess gene flow in each case (Supplementary Materials). All analysis from the Dweighted_topo statistics show generally similar putatively introgressed regions between the different analyses, although some species combinations suggested more than others (Figure S7). In both cases, very little of the genome had a higher weighting for the other alternate topology (non-species or introgression tree), as would have been indicated by Dwt < 0 (Figure 3b). In total, 52Mb of the genome was consistently found to be introgressed between *O. leucostictus* and *O. niloticus,* and 4Mb introgressed between *O. chungruensis* and *“*Chambo” (Figure 3b), from 761Mb in the *M. zebra* reference genome linkage groups. There was significantly more of the genome introgressed in both comparisons (816Kb, with regions across all 22 linkage groups; Figure 3b) than would be expected randomly (permutation test *p*=0.0002; Figure S8).

The nature of *O. chungruruensis* introgression with “Chambo” and *O. shiranus shiranus* was investigated by ADMIXTURE analysis assuming two populations (K=2; cross-validation score = 0.48), with individuals of “Chambo”, *O. chungruruensis* and *O. shiranus shiranus* from the Lake Malawi basin (*O. placidus* and *O. shiranus chilwae* were not recorded here so was not used). This suggested that 47-48% of the *O. chungruruensis* genome had affinity with “Chambo”, and 52-53% with *O. shiranus shianus*. When assuming three populations (K=3; cross-validation score =0.58), *O. chungruruensis* formed its own group (Figure 4). Diagnostic SNPs were also identified for “Chambo”, the *O. shiranus - O. placidus* group and *O. chungruruensis* (Supplementary Results). Note that diagnostic SNPs could not be detected for *O. shiranus shiranus* alone, and so the monophyletic *O. shiranus-O. placidus* group as a whole was used. These showed similar frequencies of SNPs fixed for either the reference or the homozygous alternate alleles of both “Chambo” and *O. shiranus - O. placidus*, with around half of diagnostic SNPs heterozygous (Figure 5b). Across the 22 linkage groups, 4,300 windows (each of length 200 SNPs) were found where all haplotypes *O. chungruruensis* and *O. shiranus- O. placidus* were sister taxa, compared to 2,247 windows where *O. chungruruensis* and “Chambo” were sister. Based on values of π and Dxy within these windows, we inferred introgression times of 21,520 (CI 16,374-47,075) years ago between *O. shiranus/ O. placidus* and *O. chungruruensis* and 13,565 (CI 10,321-29,673) years ago between “Chambo” and *O. chungruruensis*.

**Figure 4.**
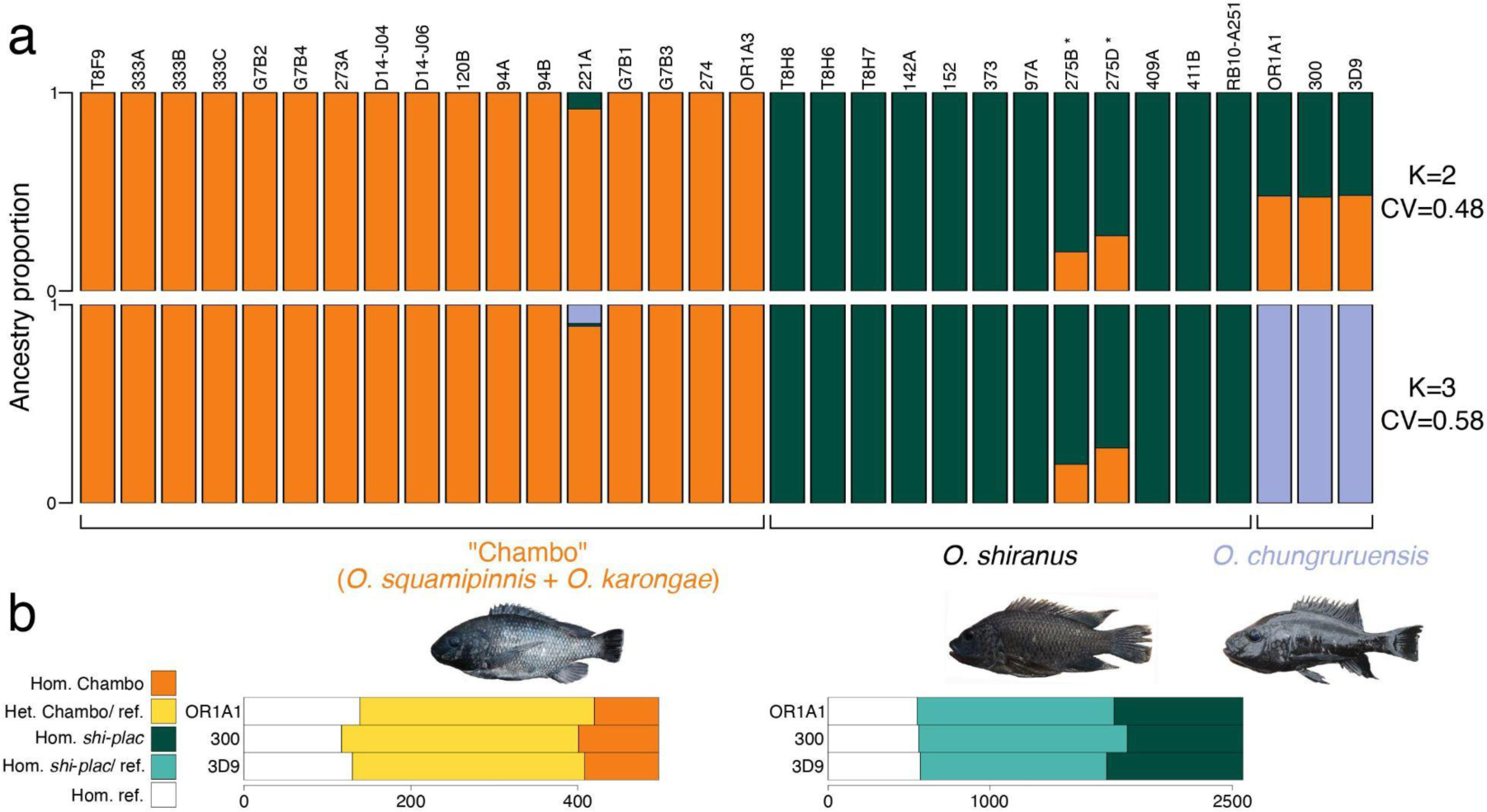
a) ADMIXTURE analysis of all *O. shiranus-O. placidus*, “Chambo” and *O. chungruruensis* at *K*=2 and *K*=3. b) The proportion of species diagnostic SNPs which are either homozygous for the alternate allele (Hom.), heterozygous (Het.), or homozygous for the reference allele (ref.) in the three *O. chungruruensis* individuals (abbreviation *shi-plac* - *O. shiranus + O. placidus*). * indicates genetically identified *O. shiranus shiranus* x “Chambo” hybrids in Lake Itamba.

**Figure 5.**
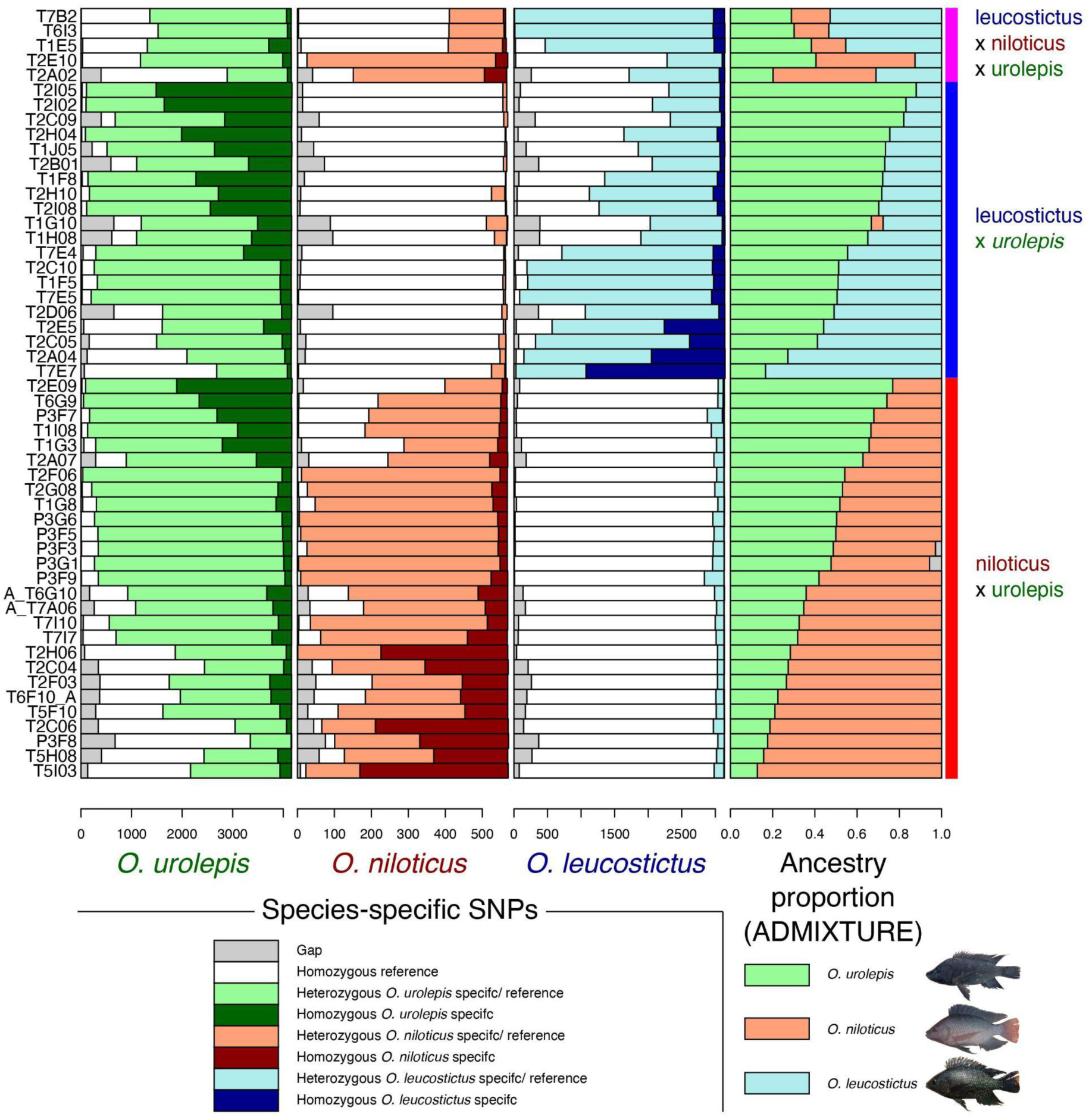
Species diagnostic SNP frequency for each individual inferred to be a hybrid between the potential parent species of *O. leucostictus*, *O. niloticus* and *O. urolepis* from ADMIXTURE analyses (ancestry components on far-right) (one per row). Diagnostic SNPs shown for *O. urolepis* (left), *O. niloticus* (mid-left), *O. leucostictus* (mid-right).

### Contemporary hybridization ongoing across Tanzania

Eleven different drainage basins and three aquaculture establishments across Tanzania were tested for the presence of hybrids (Figure 1). A total of 76 hybrids (Table 1) were identified. These were identified from across seven drainage basins: Ruvu River (30/46 tested individuals identified as hybrid), Pangani River (15/51), Rufiji River (11/56), Wami River (6/109), Lake Malawi (2/12), Lake Rukwa (2/29) and Ruvuma River (1/3) and Lake Chala (1/1). Hybrids were also identified at two aquaculture establishments: the Songea hatchery (7/50) and Mbarali Farm (1/24). It should be stressed that these numbers should not be taken as representative, as sampling was non-random and biased towards individuals thought to be hybrids on the basis of morphology.

**Table 1.**
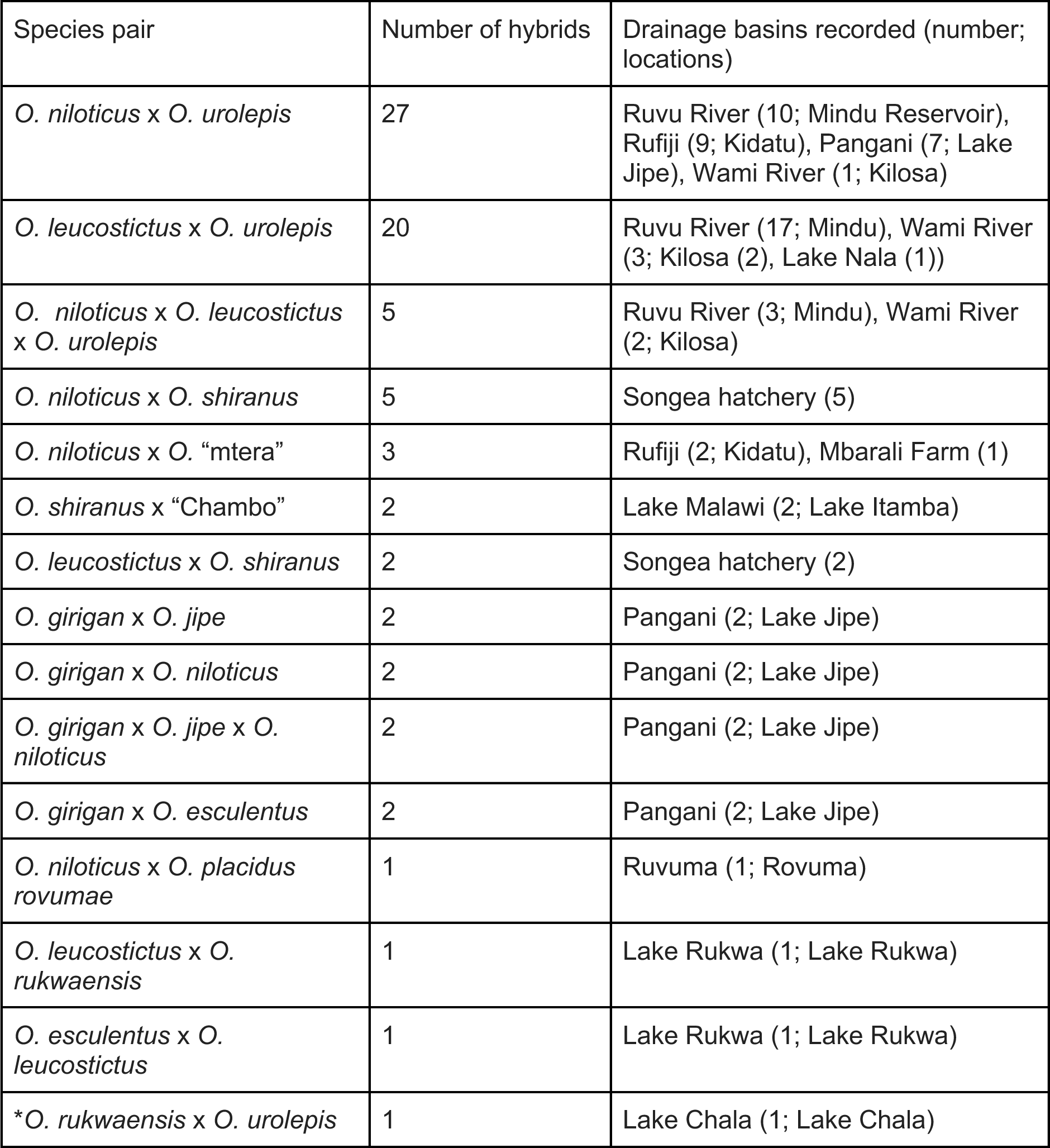
Recent hybrids identified from the supervised ADMIXTURE analysis. See Table S1 for exact locations and ancestry components of each hybrid. * indicates the individual with the “Bandia” phenotype with Lake Chala.

Most of the hybrids in our samples (68%) were between *O. urolepis, O. leucostictus* and *O. niloticus*, with none directly between *O. leucostictus* and *O. niloticus*. Species diagnostic SNPs, fixed in the confidently non-hybrid individuals of each of these three species, and with a low frequency (<0.1) in other non-hybrid individuals, were identified for each, with numbers ranging from 571 in *O. niloticus*, to 3,147 in *O. leucostictus* and 4,177 in *O. urolepis*. The hybrids identified by ADMIXTURE between the three species were investigated for these diagnostic SNPs, with results generally concordant with the ADMIXTURE ancestry components (Figure 5). The individuals with ADMIXTURE ancestry components close to 50% had high levels of heterozygosity for both species, potentially indicating F1 hybrids. When ancestry components rose above 50% for a species, it was generally associated with more homozygous species diagnostic SNPs, suggesting backcrossing.

Only 51/124 individuals morphologically suggested to be possible hybrids were genetically confirmed, ranging across the Rufiji, Ruvu River, Wami River, Zanzibar, as well as the Songea hatchery. 24 out of the remaining 321 test individuals were genetically, but not morphologically identified as hybrids. These included individuals morphologically assigned to *O. urolepis* from Mbarali Farm, the Ruvu River and the Wami River, *O. shiranus shiranus* from Lake Itamba and Songea hatchey, *O. niloticus* from Lake Jipe*, O. girigan* from Lake Jipe, *O. esculentus* from Lake Rukwa, *O. rukwaensis* from Lake Rukwa, *O. pangani* from Lake Kalimau and *O. placidus rovumae* from the Muhuwesi River (a tributary of the Ruvuma River).

The ‘Bandia’ individual, from a potentially hybrid population found in Lake Chala of unknown origin, was identified as a probable *O. rukwaensis* x *O. urolepis* hybrid, according to a panel of species diagnostic SNPs, as well as ADMIXTURE and Twisst analysis (see Supplementary Results).

## Discussion

Here, we present evidence that the genus *Oreochromis*, a group of cichlid fish important for global aquaculture, has undergone introgressive hybridization multiple times in the course of its evolution, and is now experiencing large-scale contemporary hybridization resulting from anthropogenic translocations. Almost all evidence of contemporary hybrids involved populations introduced to boost aquaculture or fisheries production, threatening the conservation of wild populations in Africa.

### Phylogenomic resolution of Oreochromis and taxonomic questions

We inferred a first phylogenomic backbone tree for the clade, building on previous work that was based on a handful of nuclear and mitochondrial markers (Ford et al., 2019). In accordance with Ford et al., (2019), we found that the *Alcolapia* group, comprised of species adapted to extreme soda lake environments (Trewavas, 1983), is nested within the *Oreochromis*, forming a clade with *O. esculentus* and *O. amphimelas*. This supports the conclusions of Ford et al. (2019) that *Alcolapia* is best considered as a subgenus within *Oreochromis*. We consistently found *O. amphimelas,* to be sister to *A. grahami.* This would suggest a single origin of tolerance of soda-lake conditions, which is present in *O. amphimelas*, albeit less extreme than *Alcolapia* (Trewavas, 1983). However, unlike Ford et al., (2019), our analyses suggests that the species previously placed in the subgenus *Nyasalapia* (“Chambo”, *O. rukwaensis*, *O. chungruruensis*, *O. variabilis*, *O. malagarasi*, *O. macrochir* and *O. karomo*) (Trewavas, 1983) do form a monophyletic clade, with the exception of *O. chungruruensis*, which has a hybrid origin between the “Chambo” and the *O. shiranus-O. placidus* group.

Our study raises several taxonomic questions related to species classifications within the *Oreochromis*. Two species of the subgenus *Nyaslapia*, *O. squamipinnis* and *O. karongae*, which are morphologically similar and endemic to Lake Malawi, comprise a single clade, but were not resolved as reciprocally monophyletic. Within the main lake, they have been differentiated largely on the basis of male breeding dress: male *O. karongae* are uniformly black (apart from white fin margins and pinkish genital papilla), while *O. squampinnis* males have a contrasting blaze of colour, generally blue, on the top of the head. In Lake Malawi itself, these traits are associated with other differences, including breeding seasonality, microhabitat and trophic morphology (Trewavas, 1983), but populations in crater lakes have not been thoroughly studied and have just been provisionally identified on the basis of male colour. Our results do not support a phylogenetic division based on male colour, particularly when the crater lake populations are considered. Ongoing admixture between species or parallel evolution may explain the evolution of these traits. The low number of specimens from Lake Malawi, as opposed to elsewhere in the catchment, prevents us from drawing clear conclusions about its populations. We were unable to sample the third *Nyasalapia* species from Lake Malawi, *O. lidole,* which has not been seen alive since 2007 and may be extinct (Konings, 2018). Genomic data on this species, potentially from museum specimens, would help with a taxonomic reassessment of the group.

Our results suggest that the current taxonomic status of *O. shiranus* and *O. placidus* needs revision. Each of these species is presently split into two geographically separated subspecies (Trewavas, 1983). Specifically, *O. shiranus shiranus* is found in Lake Malawi and the Upper-Middle Shire River, while *O. shiranus chilwae* is found in Lake Chilwa and Chiuta. Meanwhile, *O. placidus placidus* is found in the Lower Shire and Lower Zambezi, while *O. placidus ruvumae* is found in the Ruvuma system. Our results suggest that the current groupings are incorrect and should be replaced by a Lake Malawi/ Shire/ Zambezi taxon and a Chilwa/ Chiuta/ Ruvuma taxon, which more closely reflects historic and present river system connectivity. There are few, if any, morphological features supporting the current classification.

We found that the *Oreochromis* population from the Mtera Reservoir (on the Ruaha section of the Rufiji system) did not cluster with *O. rukwaensis* sampled from the neighbouring Lake Rukwa catchment. Instead, it was sister to a clade consisting of *O. rukwaensis, O. variabilis* and *O. malagarasi.* This first genomic assessment of the population is consistent with the Mtera population representing a candidate new species, which we refer to as *O.* sp “mtera”. There was some apparent introgression between *O. rukwaensis* and *O.* “mtera”, perhaps explaining the morphological similarity that led to previous assumptions of their conspecific status (Genner et al., 2018).

### Widespread ancestral introgression

We found that there was widespread, although uneven, introgression across the *Oreochromis* phylogeny. The degree of introgression was only weakly predicted by phylogenetic distance or whether species occupy the same drainage basin in the modern day. This is, however difficult to test given the highly correlated nature and lack of independence of introgression statistics, which are calculated on different subsets of four taxon across a phylogenetic tree. We may therefore have lacked power to identify these influences. We highlight two notable instances of introgression between relatively phylogenetically divergent species, primarily between the “Chambo” and *O. shiranus* (likely resulting in *O. chungruruensis*), and between *O. niloticus* and *O. leucostictus.* These latter two species coexist in their indigenous habitat of the Albertine Rift lakes Edward, George and Albert, and have been co-translocated to the same water bodies (e.g. Lake Victoria), yet are rarely found to hybridize in the modern day (Ciezarek et al., 2022; Geletu & Zhao, 2022). Our results may be reflective of gene flow between the two species at early stages of the speciation continuum (Stankowski & Ravinet, 2021).

We found that *O. chungruruensis* could have arisen as a result of hybrid speciation between “Chambo” and either *O. shiranus shiranus* or the wider *O. shiranus/ O. placidus* group. Interestingly, *O. chungruruensis* is currently classified as a member of the subgenus *Nyasalapia* alongside *O. squamipinnis* and *O. karongae* (Trewavas, 1983), which was supported by previous studies based on a handful of nuclear and mitochondrial markers (Ford et al., 2019). *O. chungruruensis* is endemic to Lake Kiungululu, an isolated crater lake within the northern sector of the Lake Malawi basin. Neither “Chambo”, or *O. shiranus* have been recorded in this lake, although “Chambo” and *O. shiranus shiranus* are found throughout the Malawi basin and co-occur in several crater lakes in the northern sector (Kingiri, Ilamba, Itamba and Ikapu). Indeed, our analysis finds two contemporary hybrids between “Chambo” and *O. shiranus shiranus* in Lake Itamba. The age of Lake Kiungulu is not known, although the nearby crater Lake Masoko has been estimated to be around 50,000 years old (Garcin et al., 2006), relatively soon before our estimated introgression ≤30,000 years ago. This may suggest a scenario where *O. chungruruensis* originated from hybridization between ancestral populations of both within the lake.

### Contemporary hybridization among Oreochromis species

Within Tanzania, there has been a long history of translocations of *Oreochromis* species for capture fisheries, dating back to at least the 1950s, in addition to ongoing aquaculture translocations, leading to the widespread colonisation by three species: *O. niloticus*, *O. leucostictus* and *O. esculentus* (Shechonge et al., 2019). Interspecific hybridization has long been recorded between *Oreochromis* species based on SNP panels, mitochondrial data and microsatellites (Geletu & Zhao, 2022), although studies based on genome-wide data are lacking. Our study has particularly demonstrated that hybridization between the invasive *O. niloticus, O. leucostictus* and the native *O. urolepis*, previously recorded in Tanzania (Ciezarek et al., 2022) is widespread. Although we found no evidence of hybrids between the invasive *O. leucosticus* and *O. niloticus*, we did find five individuals which had ancestry components of over 0.1 (10%) for both species as well as *O. urolepis*. This suggests that there may be barriers preventing *O. leucostictus* and *O. niloticus* from directly hybridizing in the modern day, despite the ancestral introgression we inferred between them. Hybridization from either species with *O. urolepis* may however mediate gene flow between them. Previous studies, based on mitochondrial data have recorded introgression directly between the two species (Ndiwa et al., 2014), although Nyingi et al. (2009) found this introgression signal was not present in the nuclear genome. Furthermore, it is not clear whether there was thorough species sampling in these studies, raising the possibility of ‘ghost’ introgression explaining these findings. Further work is necessary to determine whether there are genomic or behavioural barriers reducing the likelihood of *O. leucostictus* and *O. niloticus* hybridizing directly.

Interestingly, the Lake Chala Bandia individual appears to be a case of hybridization between two species not native to the lake (*O. urolepis* and *O. rukwaensis*), plausibly in farm ponds prior to its introduction to the lake. Previous studies on Bandia have suggested some clustering with *O. urolepis*, but with a high degree of diversity in both mitochondrial DNA and morphology (Dieleman et al., 2019; Moser et al., 2019). This is consistent with our finding of hybrid origin. Our study is however, based on only a single Bandia individual, and so more detailed population genomic analysis would be welcome.

### Concluding remarks

Several caveats need to be considered when assessing our findings. Critically, our study did not include every existing species or population of *Oreochromis* found across the globe. These missing or ‘ghost’ taxa, in addition to extinct populations of *Oreochromis* may have significantly misled analyses of ancestral introgression (Tricou et al., 2022a, 2022b). Extensive sampling of tilapia at both the species and population levels across Africa and the Middle East will be necessary to assess this, in addition to the development of bioinformatic tools which can accurately infer phylogenetic networks with large datasets, including genome-scale data and many individuals. Ghost taxa may have influenced the identification of recent hybrids, particularly in water bodies where *Oreochromis* was less thoroughly sampled. There is always the possibility for unsampled species to be responsible for the observed signals.

Our results highlight the complex relationship between farmed animals and their wild relatives. The diversity that exists within the wild may be used to enhance that of farmed populations by selective breeding (Bentsen et al., 2017). However, this potentially important diversity may also be lost due to unintentional hybridization with farm escapees. This accidental hybridization may also in turn have a harmful effect on farm populations, by potentially introducing alleles less beneficial for farming. This dynamic is particularly pronounced in *Oreochromis*, and requires ongoing genomic monitoring and assessment, to meet both food security and conservation outcomes.

## Data availability

All raw-read data generated will be made available in the European Nucleotide Archive upon publication. All SNP datasets and inferred phylogenetic trees will be made available in dryad upon publication. All custom R and python scripts will be made available on github upon publication.

## Acknowledgments

The authors declare no competing interests. Sequencing and library construction was delivered via the BBSRC National Capability in Genomics and Single Cell (BB/CCG1720/1) at Earlham Institute by members of the Genomics Pipelines Group. The computing infrastructure and data centre delivered via the NBI Research Computing are supported by the Biotechnology and Biological Sciences Research Council (BBSRC), part of UK Research and Innovation, Core Capability Grant BB/CCG1720/1.

This work was funded by BBSRC/NERC award BB/M026736/1 (GFT, MJG, FdP), and Royal Society Leverhulme Trust Africa Awards AA100023 and AA130107 (MJG, BPN, GFT, RT). WH and FDP acknowledge the support of the Biotechnology and Biological Sciences Research Council (BBSRC), part of UK Research and Innovation; this research was funded by the BBSRC Core Strategic Programme Grants BB/CSP1720/1 and its constituent work packages *-* BBS/E/T/000PR9818. AC, WH, FDP were supported through BBSRC GCRF funding (BB/P028098/1). Samples from Tanzania were collected across 2013-2016 under permit numbers 2012-393-ER-2011-103, 2014-374-ER-2011-103, 2015-83-NA-2011-103, and 2016-293-NA-2011-103. Reference samples from Uganda (Lake Albert) were collected in 2015 under permit number FS06/15/04. This study was approved by the Agricultural Research Organization Committee for Ethics in Using Experimental Animals and was carried out in compliance with the current laws governing biological research in Israel (Approval number: IL-568/15). We are grateful for the technical and logistical support from staff of the Tanzania Fisheries Research Institute. We thank Alan Smith, Steph Bradbeer, Carlos Gracida Juarez and María Lorena Romero-Martinez for assistance with sample collection and DNA extraction.

