## Supplementary Materials for "Ancient and ongoing hybridization in the *Oreochromis* cichlid fishes"

**Supplementary methods**

*Identification of individuals for backbone phylogeny*

For the 450 samples not morphologically identified as hybrid (Table S1), a neighbour-joining tree, based on nuclear distances, and a mitochondrial phylogeny were inferred. For the mitochondrial phylogeny, reads mapping against the *M. zebra* mitochondrial reference contig (NC_027944.1) in the reference genome were extracted from the mapped bam files using samtools (Li et al., 2009), and converted back to fastq using picardTools (v2.18.4; http://broadinstitute.github.io/picard/). These reads were then used to construct *de novo* mitochondrial genomes, using Norgal (v0.1) (Al-Nakeeb et al., 2017). Samples that had a mitochondrial genome of at least 10,000 base pairs assembled were rotated to have equivalent start locations (as mitochondrial genomes are circular) using MARS (Ayad & Pissis, 2017), and aligned using MAFFT (v7.271) (Katoh & Standley, 2013). A phylogenetic tree was then inferred using iqtree (v2.0), with 1,000 rapid bootstraps and automated model selection (Hoang et al., 2018; Kalyaanamoorthy et al., 2017; Minh, Schmidt, et al., 2020). For the neighbour-joining tree, genetic distances between SNPs samples were calculated using VCF2dis (https://github.com/BGI-shenzhen/VCF2Dis). These SNP-distances were converted to genetic distances by multiplying them by the number of SNPs divided by the genome size (957.5Mb). These distances were then converted according to the formula of (Dasarathy et al., 2015), which improves accuracy under incomplete lineage sorting (Rusinko & McPartlon, 2017): $d = -\frac{3}{4}log(1-\frac{4}{3}b)$, where *d* is the converted distance and *b* is the genetic distance between two individuals. These converted distances were then used to construct the neighbour-joining tree using the ‘nj’ function of the APE package within R (Paradis et al., 2004). An initial round of unsupervised per-drainage basin ADMIXTURE (Alexander et al., 2009) analysis (see “Assessment of recent hybridisation” section below) was also carried out. Reference individuals were confirmed if they were found in drainage basins where they are thought to be native, appeared monophyletic with their identified species or population in the neighbour-joining and mitochondrial trees, and showed no evidence of hybridisation in the per-drainage basin ADMIXTURE analyses. A maximum of two individuals per population per sampling location were retained.

*Backbone phylogeny reconstruction*

For the concatenation analysis, SNPs at least 10kb from any annotated gene with missing data < 10% and at least one sample with both the homozygous reference and homozygous alternate states were extracted in the 91 individuals, and lightly pruned for linkage (removing SNPs with r2 > 0.9 over 20kb windows) using bcftools (v1.12). SNPs from the mitochondrial genome and the repeat-rich LG3 mega-chromosome in the *O. niloticus* mapping were removed. A phylogenetic tree was then inferred using iqtree, with automated model detection, 5 independent runs and 1000 rapid bootstraps.

For the multispecies coalescent tree, fasta alignments were generated for the 91 reference individuals from the bam files using ANGSD (v0.923). The base with the highest depth per site was called, with a minimum quality and mapping quality of 30 and a minimum depth of 3. These alignments were split into 10kb windows using bedtools (v2.28.0). The resulting alignments were trimmed to remove columns with at least 30% gaps and individual sequences with at least 50% gaps. Alignments with a minimum length after filtering of 600bp with at least 10 retained individuals were kept. The Phi test was then used to assess each 10kb alignment for evidence of intra-locus recombination. Those without evidence of intra-locus recombination (un-corrected p>0.05) were retained, as long as the sequence immediately before them was not also retained (to avoid pseudo-replication). A phylogenetic tree was inferred for each of the retained 10kb windows using iqtree (v1.6.12), with automated model detection, 10 independent runs, 1,000 rapid bootstraps as well as 1,000 bootstrap replicates for the Shimodaira-Hasegawa (SH)-like approximate likelihood-ratio test. Outlier trips were pruned from the inferred trees using treeshrink (v1.3.1), with a quantile threshold of 0.05. Nodes with a SH-like alrt <1 (Simmons & Gatesy, 2021) or a bootstrap support < 10% were collapsed into hard polytomies before species-tree inference using ASTRAL. Four runs were carried out: one outputting each alternate quartet score, one outputting the posterior probability and one with bootstrapping with gene-tree resampling. A further run was carried out for plotting and inference of terminal branch lengths, with forced monophyly individuals from the same species. In cases where species were not monophyletic, individuals from monophyletic populations within the species were clustered together. Groups of species which formed an indistinguishable clade (e.g. *O. squamipinnis* + *O. karongae*, hereafter termed 'Chambo') were grouped together. *O. korogwe* from Nambawala were considered a separate population to *O. korogwe* from Mlinango, based on recent findings of population segregation, and possible introgression of *O. niloticus* into the Nambawala population (Blackwell et al., 2020). Robinson-Foulds (RF) distances (Robinson & Foulds, 1981) were calculated between each 10kb-window tree and the inferred ASTRAL species tree using the ETE 3 python package (Huerta-Cepas et al., 2016), in order to further quantify how widespread phylogenetic discordance was. This was carried out both using all individuals from all species (individual-level), comparing against the full ASTRAL tree, or using one individual from each species (species-level), against the ASTRAL tree with forced monophyly of species. A histogram of these distances was plotted using R. Differences between individual level and species-level were assessed by a one-way anova test.

*Assessment of ancestral introgression*

To calculate D statistics, all non-reference samples were pruned  from the SNP set and sites with at least 1 SNP in the remaining 91 individuals, with less than 50% missing data were retained. Dsuite (v0.4) (Malinsky et al., 2021) was used to calculate D statistics for each trio of species, using a jackknife block size of 1,000 SNPs and the backbone tree inferred by ASTRAL, rooted with *M. zebra*. Monophyletic species were specified as a single species in the population mapping file, whereas non-monophyletic species were split into monophyletic populations. Significance of D statistics was assessed using bon-feronni corrected block jackknife $p<0.05$. The *f*-branch metric (Malinsky et al., 2018) was also calculated, which disentangles correlated *f*4 statistics to assign gene flow to specific branches of a phylogeny. To quantify the extent of introgression, the Dp statistic (Dp = $\frac{ABBA - BABA}{ABBA + BABA + BBAA}$) was also calculated (Hamlin et al., 2020) and added to the Dsuite output using a custom python script. The average Dp per unique combination of P2 and P3 was calculated, and Dp values from the *M. zebra* and *O. niloticus* mappings were compared by Pearsons correlation.

One disadvantage of site-pattern based ABBA-BABA (D) tests, is that as evolutionary time between populations increases, so does the likelihood of model violations. Particularly, it is likely that back-mutations occur, and the rate of evolution varies between populations, which may cause false positive tests. We therefore also carried out tests based on gene-tree discordance, utilising the discordant count test (DCT) and branch-length tests (BLT) of (Suvorov et al., 2022). The ASTRAL tree was used as the backbone, and the individual 10kb-window trees used for its inference were used for discordance analysis, all rooted with *M. zebra*. A Ddct statistic, analogous to the ABBA-BABA D statistic was calculated as follows, where disc1 and disc2 are the counts of the two discordant topologies for each rooted triplet. $Ddct = \frac{disc1 - disc2}{disc1 + disc2}$. Significance was assigned for a trio if both the BLT and DCT had bon-feronni corrected $p < 0.05$ (chi-squared for DCT, Mann-Whitney u for the BLT). Gene-tree discordance at each node of the ASTRAL phylogeny was also calculated for both mapping datasets. All gene-trees, rooted with *M. zebra* using Newick Utilities (v1.5), were compared against the phylogeny using iq-tree (v2.0). For each node, chi-squared was used to assess whether the frequencies of the discordant topologies significantly differed (bon-feroni corrected $p<=0.05$) from each other. This possibly indicates introgression, as under incomplete lineage sorting alone frequencies would be expected to be equal (Minh, Hahn, et al., 2020).

*Analysis of putative introgression events*

For the Twisst analysis (see main text), individuals from the relevant populations were extracted from the original filtered vcf files (just the outgroup *M. zebra* mapping), with only sites with at least 3 SNPs in the target populations, less than 10% missing taxa retained, and located on the main 22 linkage groups. Target populations were selected to include only four populations in each comparison, as the number of topologies to compare increases exponentially with increasing populations. The two putatively introgressed populations, in addition to one population sister to each in the species tree were selected. The weightings of the topology matching the species tree were then compared with those of the two alternative topologies, one of which matches the putative introgression event, across the genome. To test for an effect of the sister population used, three iterations were carried out with a different sister population used in each. A final run was then carried out with the sister populations grouped together into one pseudo-population. These extracted genotypes were then phased and imputed using beagle (v4.1), with a window size of 10,000 bp and overlap of 1,000 bp. These were then converted to the geno format using genomics_general. Phylogenetic trees were inferred along the length of the linkage groups in sliding windows of 200bp, with 40bp overlap, using iqtree (v1.6.12), and automated model detection with ascertainment-bias correction. In order to visualise regions where each of the three topologies was most prevalent, a custom statistic was plotted along the length of the linkage groups. In the following equation *sptree* refers to the weighting of the species tree, *disc1* the weighting of the putative introgression tree, and *disc2* the weighting of the other possible topology. All windows where there was Dwt of exactly 1.0 (full support for the introgressed phylogeny) over a span of at least 50kb were extracted, using a custom python script, and windows where this was consistently the case across the four different species subsets were extracted using BEDTools (v2.30.0) (Quinlan & Hall, 2010) intersect, sort and merge. From these final, merged putative introgressed region bed files, it was then assessed whether there were more overlapping regions between introgression events than expected, using a permutation test, carried out in the 'regioneR' R package (Gel et al., 2016), using the overlapPermTest function, with 5,000 iterations and the randomizeRegions randomisation function.

We then assessed cases where Twisst showed no clear excess of either the species or introgression trees for evidence of hybrid speciation. We carried out ADMIXTURE analysis for all individuals of the three species (the two putative parent species and the putative hybrid species) at both K=2 and K=3, comparing the ancestry components and cross-validation scores of each. Diagnostic SNPs were identified for the three species, by first filtering any putative hybrids from the full *M. zebra* mapping SNP set and finding SNPs in any of the three species, and removing those with any missing data for the three target species. From this reduced set, SNPs unique to the possible hybrid species, identified from ancestral introgression analyses as well as ADMIXTURE output, were identified, using bcftools view with "-x" for private alleles were extracted, and sites where the diagnostic SNP had an allele frequency of at least 0.75 were counted. The putative hybrid species was then filtered from the SNPs set, and sites which segregated the two parent species (with diagnostic allele frequency > 0.75) were extracted. For each individual of the putative hybrid species, it was counted how many of these diagnostic SNP were heterozygous, homozygous reference, or homozygous alternate (i.e. the species diagnostic site was present and homozygous). Under a situation of hybrid speciation, it may be expected that the hybrid ancestry has ADMIXTURE ancestry components of roughly 50% for each of the parent species, has its own diagnostic SNPs (suggesting it is not a very recent hybrid), and roughly equal proportions of fixed SNPs from either parental species. Recent hybrids would be mostly heterozygous sites from each species, and backcrosses would have an excess ancestry component and proportion of fixed SNPs for one species. In order to putatively date when introgression events occurred, phylogenetic trees were inferred across the 22 long linkage groups in 200 SNP windows, with no overlap, using all individuals from the putative hybrid species and its two parents, as well as the outgroup *M. zebra* for rooting. Input phased geno format files were prepared as with the Twisst analysis described earlier. Regions where the putative hybrid species was monophyletic with either parent species were extracted. Dxy between each population, and π within each population, were calculated within these windows, using the popgenWindows.py script within genomics_general. In order to calculate the number of callable sites within each window, which is necessary to get accurate estimations of Dxy and π, which require invariant as well as variant sites, the number of sites which had total depth within the limits filtered for in our SNP calling pipeline were calculated using samtools, and used to adjust Dxy and π values. To get an estimate of the divergence dates between pairs of population, the average pi value for either population was subtracted from the Dxy between the two populations (Shang et al., 2023). This was then divided by an estimated cichlid mutation rate of $3.5 \times 10^{-9}$ (confidence interval [CI] $1.6\text{-}4.9 \times 10^{-9}$) substitutions per bp per generation (Malinsky et al., 2018), which was assumed to be one year (Blackwell et al., 2020).

*Assessment of recent hybridisation*

Following identification of recent hybrids (see main text), we further investigated the hybridisation involving *O. urolepis*, *O. leucostictus* and *O. niloticus*. We first identified segregating SNPs unique to each of the three species. A vcf file was extracted with SNPs found in individuals confidently not characterised as hybrid (those that had only one ADMIXTURE ancestry component of at least 0.001 and did not share a sampling site with any identified hybrids) were extracted, using bcftools view. SNPs fixed in each of the three species, with a low frequency (<0.1) across all other species were then extracted using a custom python script. These were considered the species-diagnostic SNPs, although we note that they are still present at a low frequency across the *Oreochromis* radiation. All of the hybrids between the three species inferred from ADMIXTURE analysis were screened against these species-specific SNP sets to record how many were fixed for the species-specific alleles, fixed for the reference allele or heterozygous. We would expect first generation hybrids to be heterozygous for most of the species-specific SNPs, with increasing amounts of fixed species-specific SNPs with backcrossing.

We also specifically analysed the potential hybrid origin of a ‘Bandia’ individual from lake Chala. This fish is hypothesised to be stocked from elsewhere in Tanzania may be an invasive population of *O. korogwe* (it has previously been labelled as *O. cf korogwe*), Previous mitochondrial studies have suggested a high degree of genetic variability, with many individuals closely related to *O. urolepis* and no evidence of ongoing *O. hunteri*, although a hybrid origin if the Bandia is possible (Dieleman et al., 2019). We identified species specific SNPs for each species in the 91 individual reference dataset, and counted how many species-specific alternate alleles were present in the Bandia individual. Private, species specific SNPs were identified using bcftools view, with only SNPs fixed in each species counted as species-specific. Ancestry for all species where the Bandia individual had an alternate allele for at least 10% of the species-specific SNPs were further investigated using ADMIXTURE, using all reference individuals from these species as well as the Bandia individual (filtering SNPs as above), using k at all values between 1 and the number of tested species plus two. We also carried out TWISST analysis (Martin & Van Belleghem, 2017) to assess phylogenetic relationships across the genome, utilising all relevant species, and took the average weighting where the Bandia individual was sister to any of the other species. For this, we first phased relevant genotypes, using beagle (v4.1) (Browning & Browning, 2011), with window size of 10,000 bp and overlap of 1,000 bp. These were then converted to the geno format using genomics_general. Phylogenetic trees were inferred along the length of the linkage groups in non-overlapping sliding windows of 200 SNPs, using iqtree (v1.6.12), and automated model detection with ascertainment-bias correction.

**Supplementary results**

*Sampling, sequencing, read mapping and SNP calling of 23 species against O. niloticus reference*

Reads from each of the 575 individuals were mapped against the *O. niloticus* reference, showing a slightly higher mapping percentage than against *M. zebra* (see main text), with an average depth of 7 (range 1.8-20.1) and average paired mapping of 90% (range 41-97%; table S1). A total of 68,783,458 filtered SNPs were called against the *O. niloticus* genome.

*Phylogenetic inference*

A total of 1,445,653 non-coding SNPs were used for the *O. niloticus* mapping maximum-likelihood tree, and 14,744 recombination free 10kb windows were used for the *O. niloticus* mapping ASTRAL tree. As with the *M. zebra* mapping, there was significant Robinson-Foulds (RF) discordance between each 10kb window and the species tree at both the individual (none less than 0.38) and population (none less than 0.11) levels, with no significant difference in RF between the individual and population levels ($p$=0.5). Results in the ASTRAL analyses were identical with the *M. zebra* mapping at the population level, with minor differences in the ML analysis (Figure S2-5).

*Widespread ancestral introgression*

Consistently, we identified a wide degree of ancestral introgression. Out of these 40,849 significant trios for both the blt and dct tests for the *M. zebra* mapping, there were 3,194 pairs of individuals with evidence of introgression out of a possible 4,095 (i.e. in the 4 taxa tests ((ind1, ind2), ind3), outgroup); testing for introgression between ind1 and ind3, there were 3,194 unique combinations of ind1 and ind3 - each significant with multiple tests with different individuals as ind2, involving 326 unique species pairs, out of a possible 406. For the *O. niloticus* mapping dataset, 47,937 out of the 113,562 were significant for both the blt and dct tests, involving 3,324 unique pairs of individuals, out of a possible 4,095 and 334 unique pairs of species, out of a possible 406. There were 322 unique pairs of species with evidence of introgression in both the *M. zebra* and *O. niloticus* mapping datasets; 5 only in the *M. zebra* only dataset and 12 only in the *O. niloticus* mapping dataset. In the *O. niloticus* mapping, 10kb window-tree concordance factors different from expectations under incomplete lineage sorting (ILS) in 43/89 nodes.

D statistics suggested significant introgression in 2,558 out of the 2,925 in the *O. niloticus* dataset (Table S2). 2,433 trios were significant in both the *M. zebra* and *O. niloticus* mapping datasets; 225 were only in the *M. zebra* dataset, and 125 only in the *O. niloticus* dataset. Dp values, showing the extent of introgression between populations, were tightly correlated (r=0.97) between the *M. zebra* and *O. niloticus* mapping datasets (Figure 2; table S2). F-branch statistics for the same comparisons in the *M. zebra* and *O. niloticus* were tightly correlated ($r$=0.97,$p$<2.2e-16; Figure S6c). Four separate Twisst analyses were carried out to assess introgression between *O. niloticus* and *O. leucostictus*. with each containing the reference individuals for both species as well as those for i) *A. grahami* and *O. aureus*; ii) *O. esculentus* and *O. spilurus*; iii) *O. variabilis* and *O. aureus*; and iv) a combination of all the previous species, with *A. grahami*, *O. esculentus* and *O. variabilis* as one group and *O. aureus* and *O. spilurus* as the other. Similarly, four analyses were carried out with *O. karongae*, *O. squamipinnis*, *O. chungruruensis* and i); *O. korogwe* (from Milingano) and *O. macrochir* ii); *O. rukwaensis* and *O. placidus rovumae* iii) *O. variabilis* and *O. mossambicus* and iv) a combination of all these species, with *O. mossambicus*, *O. placidus rovumae* and *O. korogwe* as one group and *O. rukwaensis*, *O. variabilis* and *O. macrochir* the other. These species and comparison were selected so there was one species sister to either of the target species, and no obvious confounding pattern of gene flow.

In order to identify diagnostic SNPs for the “Chambo”, *O. shiranus/ placidus* group or *O. chungruruensis*, SNPs found in any of the target populations were identified from the full *M. zebra* mapping dataset, after the exclusion of any morphologically identified hybrid, individual of the target species putatively identified as hybrid, or *O. urolepis*, given the wide extent of gene flow with the *shiranus/ placidus* group indicated by *f*-branch (Figure S6). From this dataset, 38 *O. chungruensis* - specific SNPs were identified as well as 497 “Chambo” and 2,565 *shiranus/ placidus* diagnostic SNPs. The three *O. chungruruensis* individuals were 56-57% heterozygous for the chambo-specific SNPs, 24-28% homozygous reference and

16-20% homozygous alternative. For the *shiranus/ placidus* diagnostic SNPs, they were 45-50% heterozygous, 21-22% homozygous reference and 28-33% homozygous reference

*Genetic identification of the 'Bandia' individual*.

To test the unknown origin of the Bandia individual, we tested it against a panel of species diagnostic SNPs. Three species were found for which the Bandia individual had at least 10% of the tested SNPs; *O. korogwe* (16% of 75), *O. rukwaensis* (30% of 791) and *O. urolepis* (90% of 755). The Bandia had previously been hypothesised to be of *O. jipe* or *O. hunteri* origin, but only one species-specific SNP of *O. hunteri* was found (not in the Bandia individual), and only 1% of the 7,313 *O. jipe* specific SNPs were found. These species were added to the ADMIXTURE analysis anyway due to previous hypotheses about its origin, as well as the closely related *O. girigan* (5% of 38,686). ADMIXTURE analysis indicated that it had an ancestry component of 75% for *O. urolepis* and 25% for *O. rukwaensis*. Topology weighting analysis indicated that the Bandia individual is most closely related to *O. rukwaensis* (49.7% of the total weighting) or *O. urolepis* (27.1%) in most of the genome, with *O. korogwe* (4.6%), *O. girigan* (3%), *O. jipe* (2.8%) and *O. hunteri* (2.3%), been infrequently sister to Bandia. Together, this suggests that the Bandia individual is an *O. rukwaensis* x *O. urolepis* hybrid.

**Supplementary Figures**

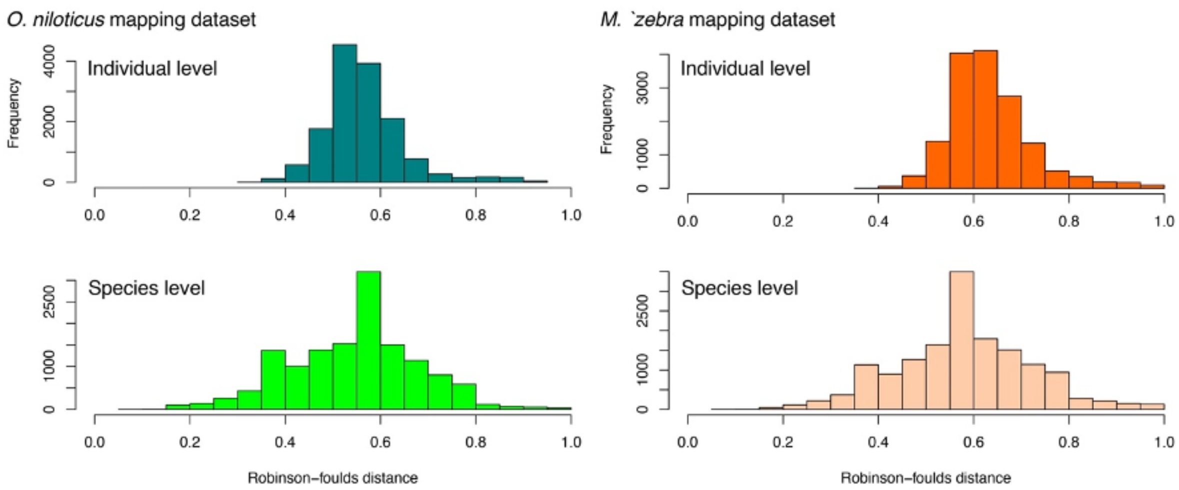


Figure S1. Robinson-foulds distances between each 10kb window tree and the species tree for the *O. niloticus* (left) and *M. zebra* (right) mappings, at both the full individual (top), and species (bottom) levels.

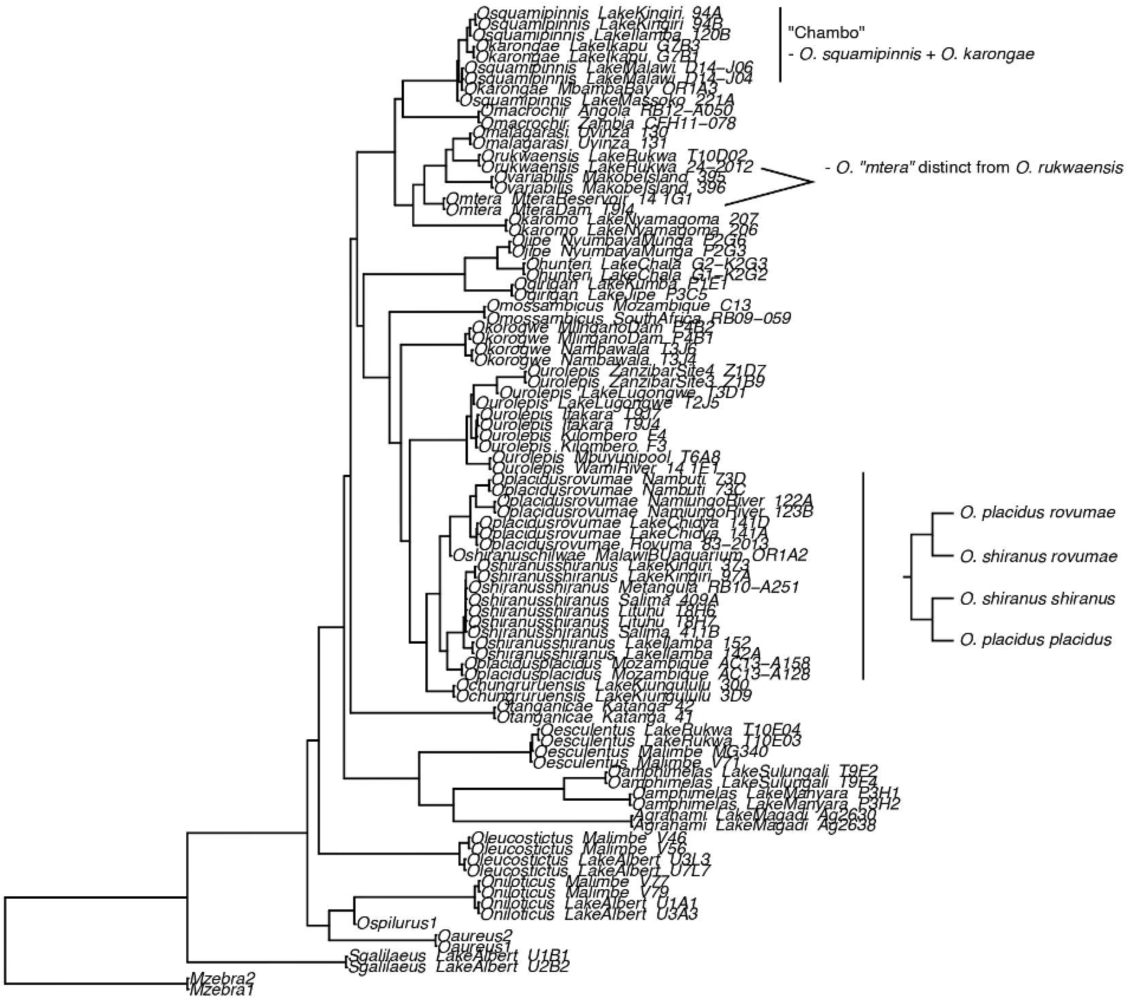


Figure S2. Multi-species coalescent phylogeny inferred using ASTRAL from 10kb windows mapping against the *M. zebra* reference genome. The “Chambo” group, whereby *O. squamipinnis* and *O. karongae* do not segregate, putatively novel species *O. “*mtera*”*, and lack of segregation between *shiranus* and *placidus* are highlighted.

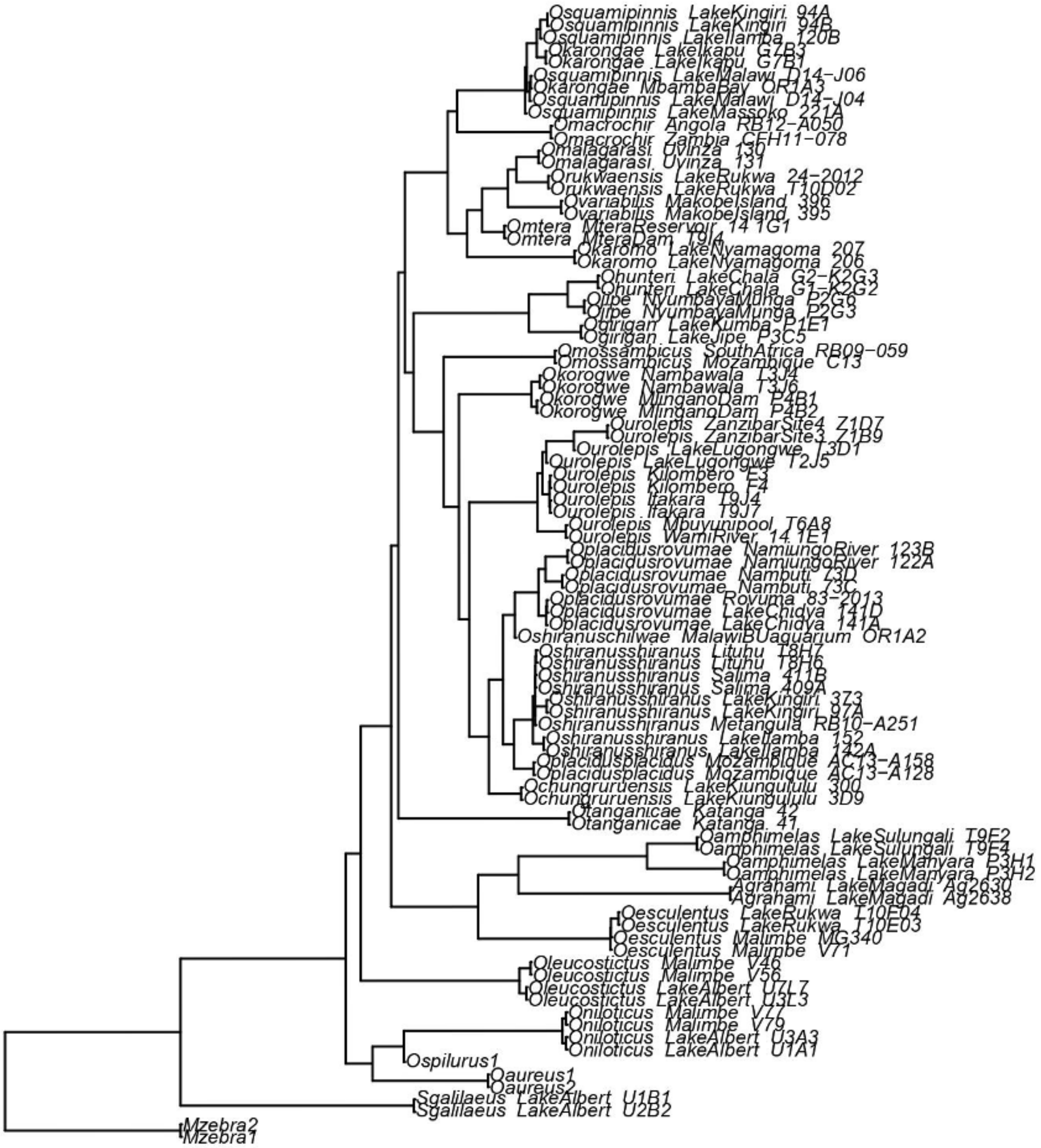


Figure S3. Multi-species coalescent phylogeny inferred using ASTRAL from 10kb windows mapping against the *O. niloticus* reference genome.

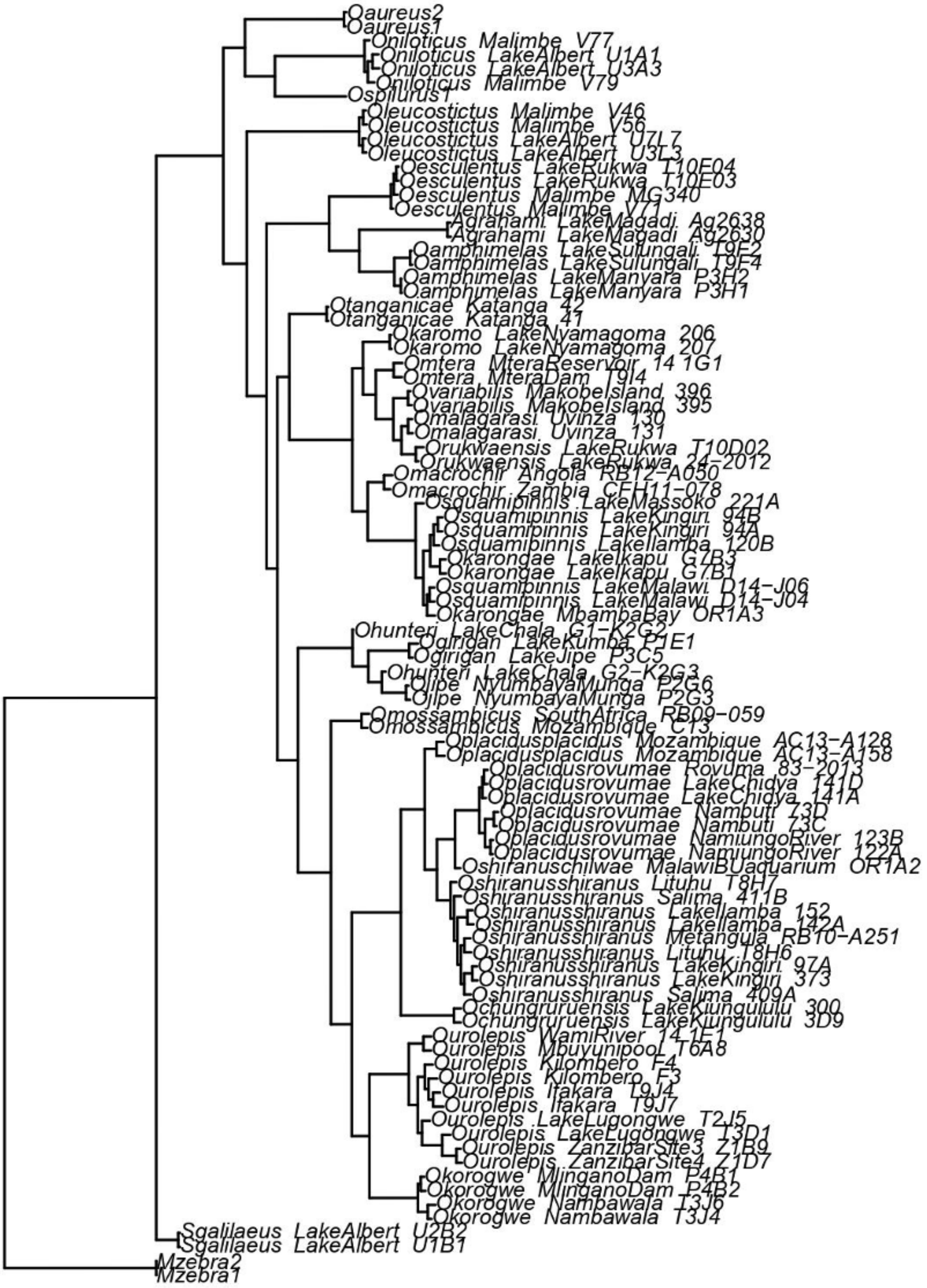


Figure S4. Maximum likelihood Phylogeny inferred using iqtree from genome-wide SNPs called against the *M. zebra* reference genome.

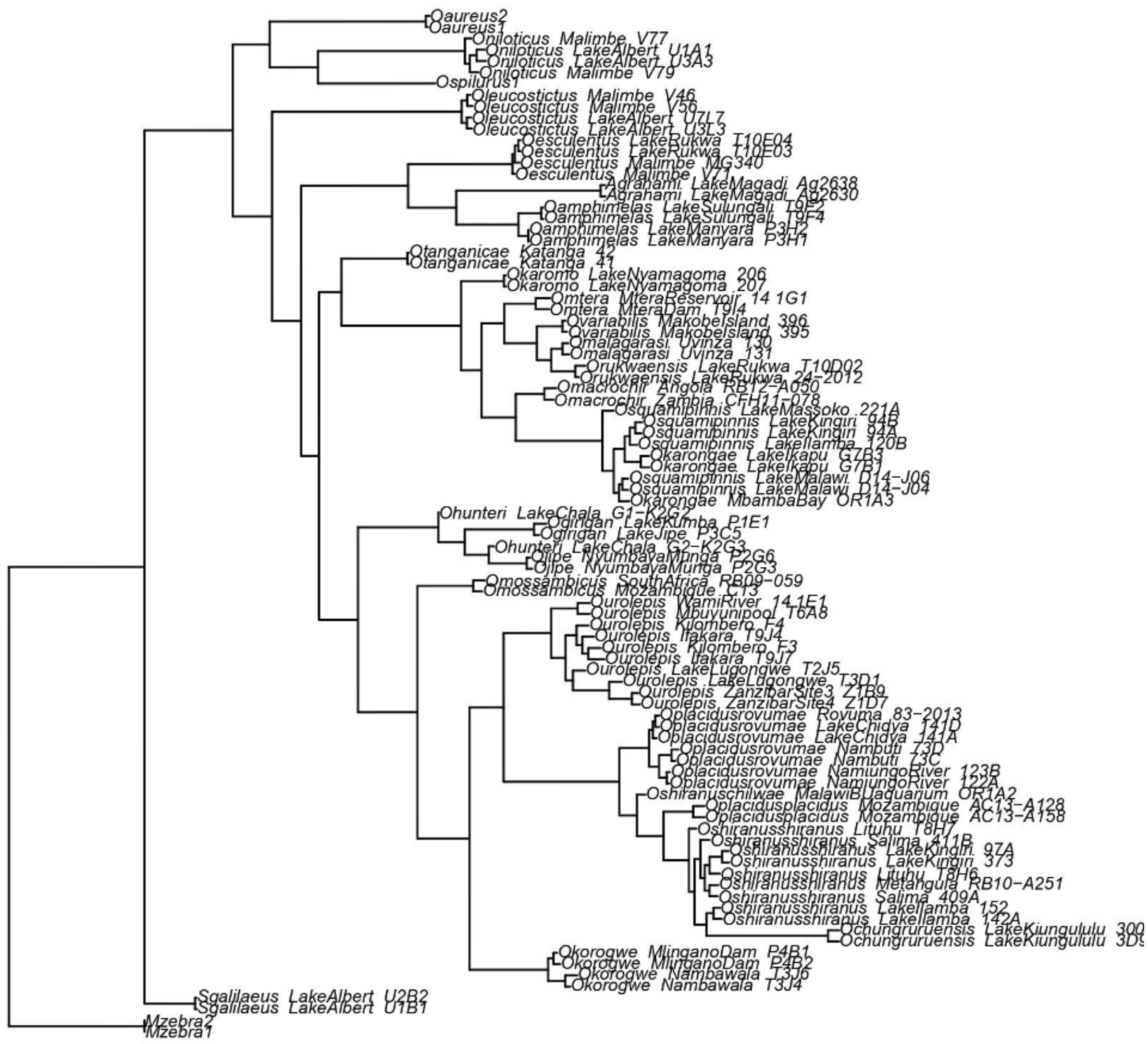


Figure S5. Maximum likelihood Phylogeny inferred using iqtree from genome-wide SNPs called against the *O. niloticus* reference genome.

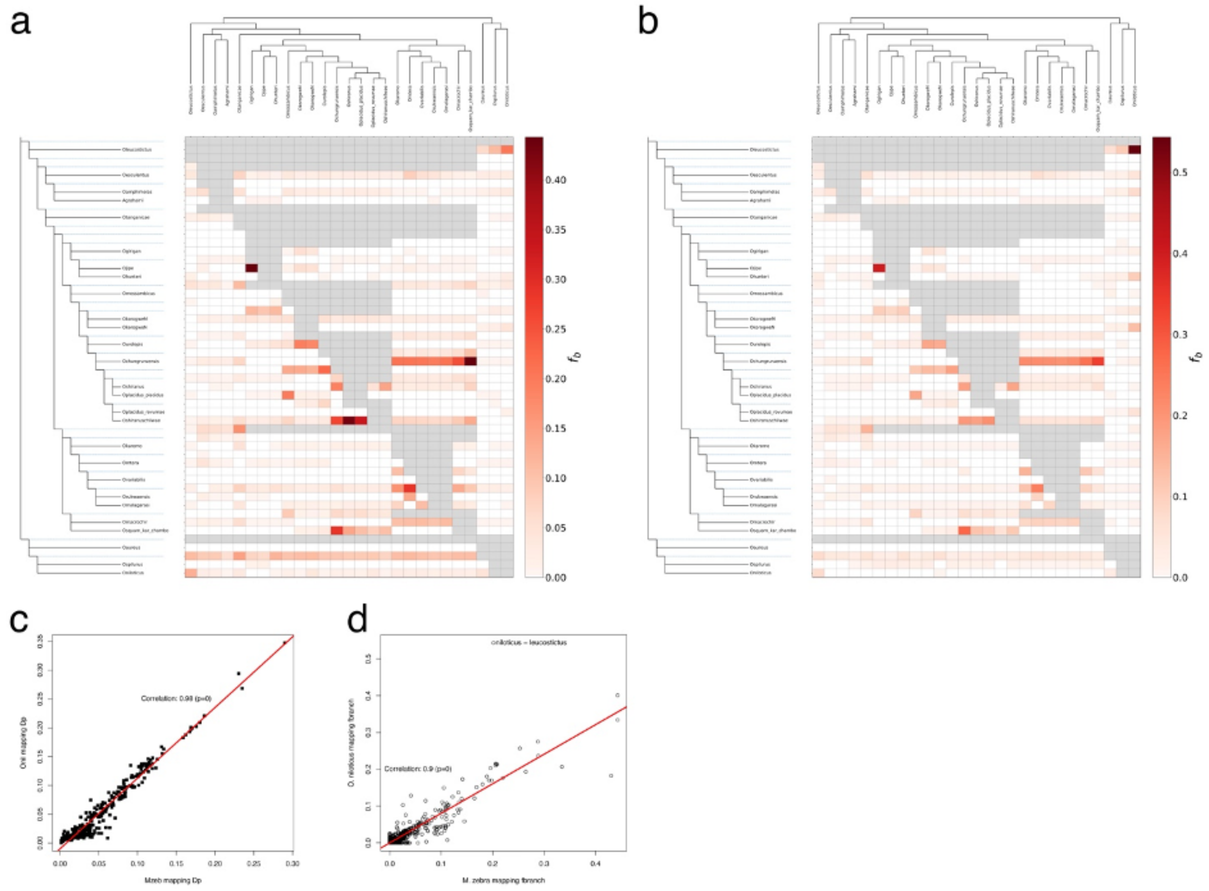


Figure S6 a) F-branch heatmap mapped from the *M. zebra* mapping; b) F-branch heatmap from the *O. niloticus* mapping; c) Correlation in Dp values from the *M. zebra* and *O. niloticus* mappings; d) Correlation between Fbranch values from the *M. zebra* and *O. niloticus* mapping datasets.

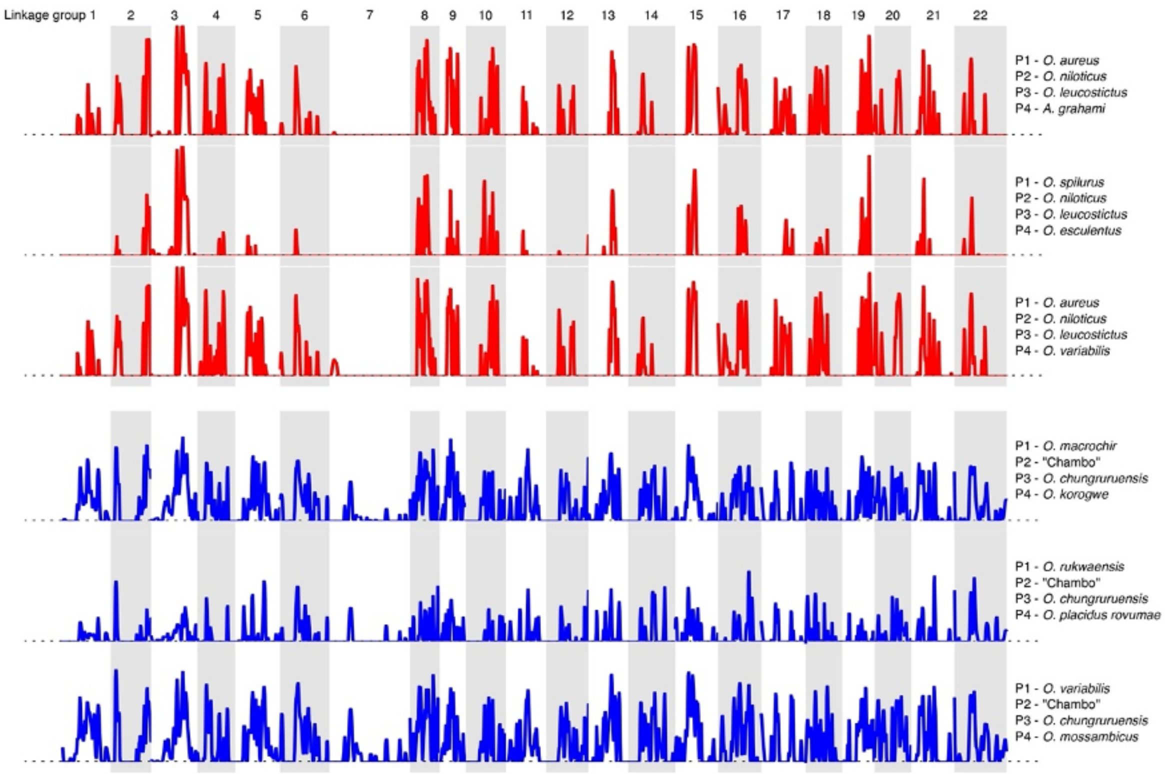


Figure S7. Comparison of Dwt values showing putative introgression analyses for the three iterations of *O. leucostictus* <-> *O. niloticus* analysis (top three rows; red, four taxon used in each comparison shown on the left), and "Chambo" <-> *O. chungruruensis* analysis (bottom three rows; blue).

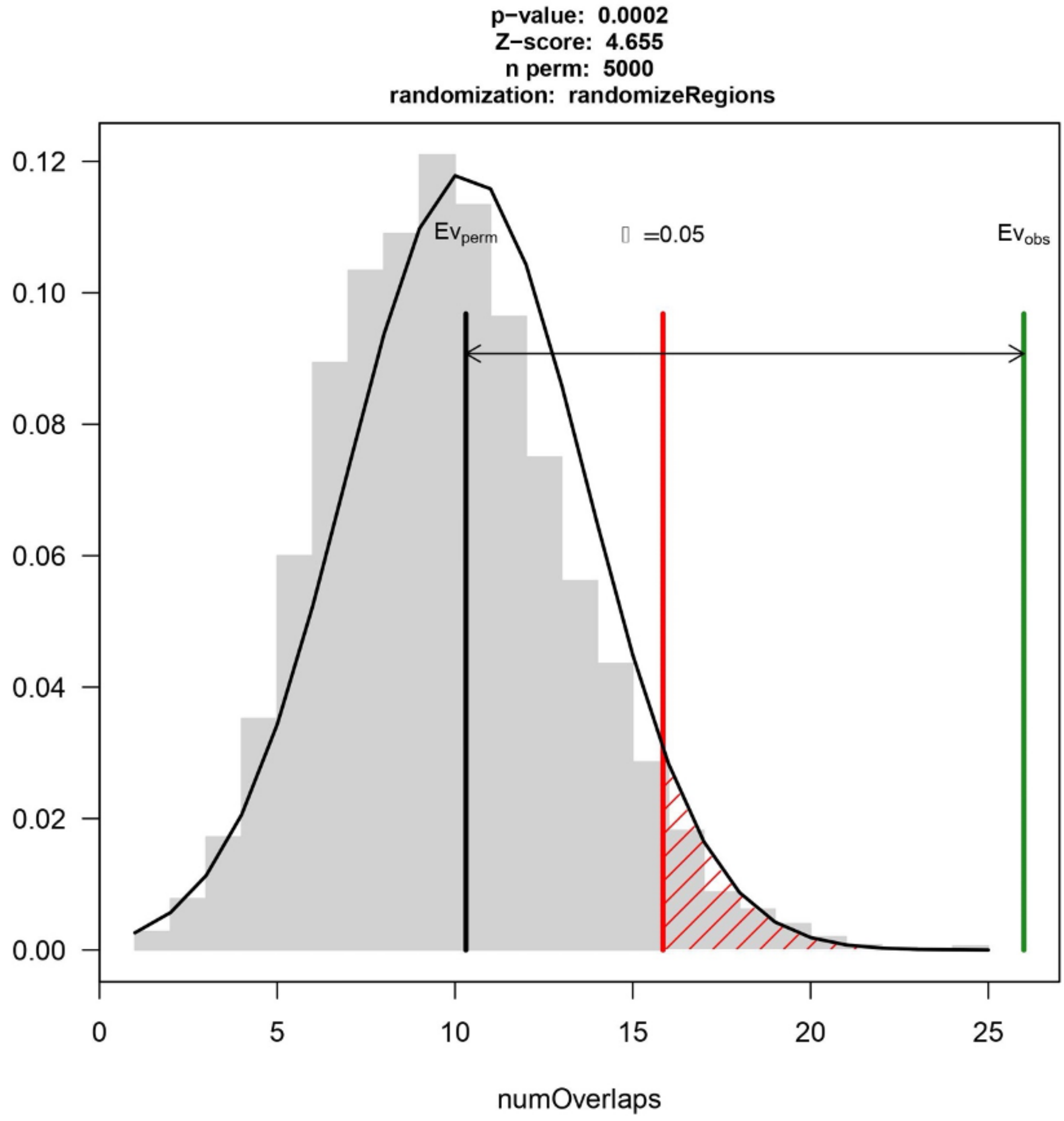


Figure S8. Permutation test analysing overlapping regions between the *O. chungruruensis* <-> “Chambo” and *O. niloticus* <-> *O. leucostictus* introgression events, output by regioneR package.
